# Allostery through DNA drives phenotype switching

**DOI:** 10.1101/2020.07.04.187450

**Authors:** Gabriel Rosenblum, Nadav Elad, Haim Rozenberg, Felix Wiggers, Hagen Hofmann

**Affiliations:** Department of Structural Biology, Weizmann Institute of Science, Herzl St. 234, 76100 Rehovot, Israel; Department of Chemical Research Support, Weizmann Institute of Science, Herzl St. 234, 76100 Rehovot, Israel

## Abstract

Allostery is a pervasive principle to regulate protein function. Here, we show that DNA also transmits allosteric signals over long distances to boost the binding cooperativity of transcription factors. Phenotype switching in *Bacillus subtilis* requires an all-or-none promoter binding of multiple ComK proteins. Using single-molecule FRET, we find that ComK-binding at one promoter site increases affinity at a distant site. Cryo-EM structures of the complex between ComK and its promoter demonstrate that this coupling is due to mechanical forces that alter DNA curvature. Modifications of the spacer between sites tune cooperativity and show how to control allostery, which paves new ways to design the dynamic properties of genetic circuits.

---

Allostery is the structural coupling between ligand sites in biomolecules. Binding of a ligand to one site facilitates or hampers the binding of a second ligand to a distant site^1-5^. The resulting cooperativity regulates the activity of many proteins and molecular machines^6-8^ but it is also key for the behaviour of genetic circuits with binary^9^, oscillatory^10^, excitable^11,12^, or pulsing^13^ dynamics. The past decades have seen growing evidence that allostery is also an inherent property of DNA^14-25^, which has far reaching consequences for our understanding of promoter sequences. Yet, most insights on DNA-mediated allostery upon transcription factor (TF) binding were either based on artificial promoters^14,26^ or found to be short-ranged^19,20,25^. Whether natural promoters evolved to efficiently transmit allosteric signals across many nanometres remained largely unclear. Here, we show that *Bacillus subtilis* bacteria utilize long-range allostery in a stochastic and reversible phenotype switch (Fig. 1a). In the competent phenotype, *B. subtilis* can take up DNA from the medium^11,27^. The master regulator of the switch from vegetative to competent cells is the TF ComK^11,12^. The model postulates a positive feedback regulation of *comK* gene expression once ComK levels stochastically surpass a critical threshold (Fig. 1b)^28^. The critical threshold acts like an analog-to-digital converter: the ComK target promoter is inactive at low concentrations but it switches cooperatively to an active state within a narrow ComK concentration range (Fig. 1c)^11,12^. Yet, how the promoter mediates cooperative ComK binding is not known. By integrating single-molecule Förster resonance energy transfer (smFRET) and cryo-electron microscopy (cryo-EM) we show how ComK binding at one promoter site enhances binding to a distant site via allosteric changes of DNA structure.

**Fig. 1.**
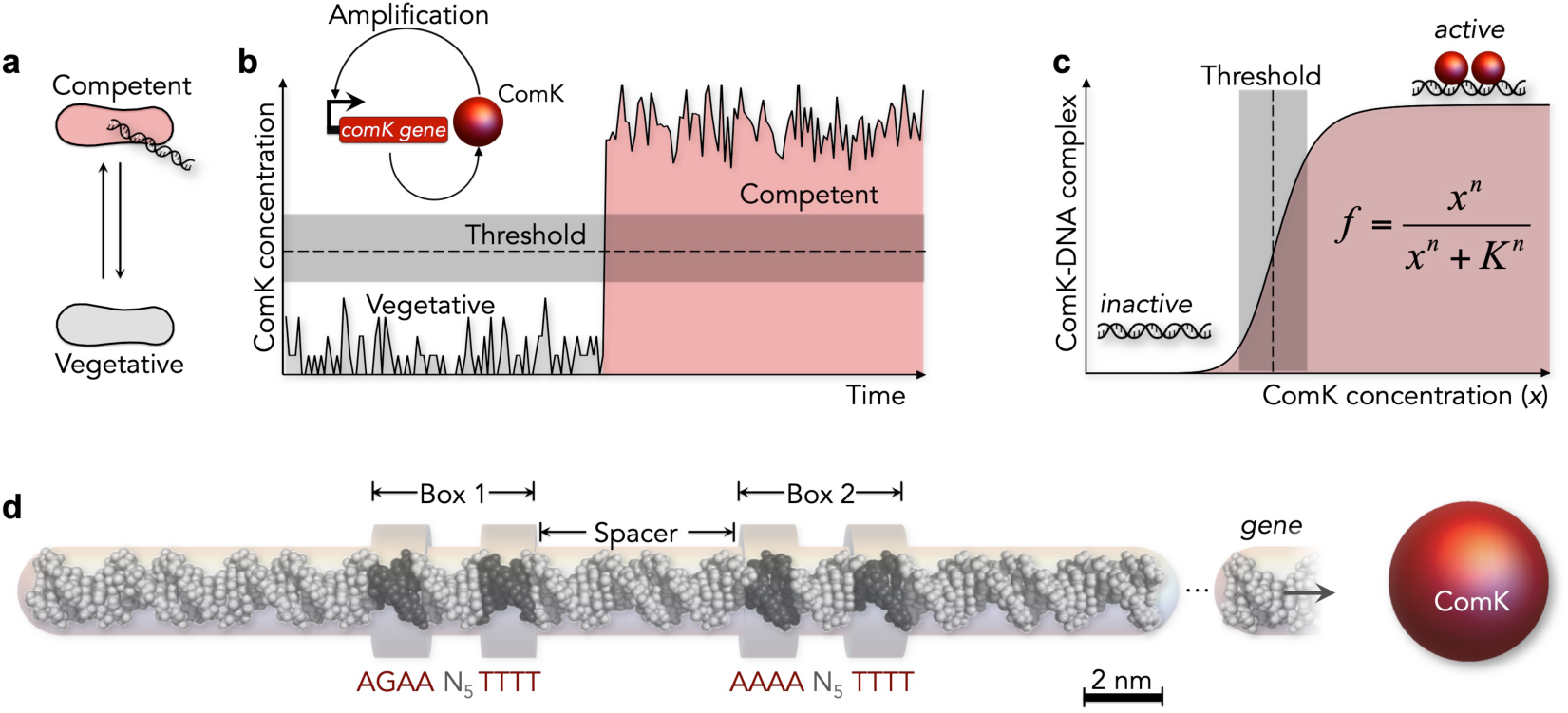
Schematics describing the model for phenotype switching in *B. subtilis*. (**a**) *B. subtilis* can reversibly switch to a competent phenotype. (**b**) Switching is triggered by copy number fluctuations of ComK (gray) that, once stochastically exceeding a threshold (dashed line), cause its auto-amplification to high concentrations (red). (**c**) The threshold requires a cooperative DNA-binding of ComK, modelled by the Hill equation with cooperativity *n*, affinity *K*, and fraction of complexes *f*. (**d**) ComK promoters consist of two boxes separated by a spacer. Here, the *comG* promoter with an 18 bp spacer is shown. For size comparison, a ComK monomer (Stokes radius: 2.3 nm, Extended Data Fig. 2) is shown as red sphere.

## Distant binding sites communicate

ComK target promoters consist of two elements (box 1 and box 2 hereafter) separated by spacer sequences of variable length (Fig. 1d)^30^. Three spacers of 8, 18, and 31 base pairs (bp) are known in the ComK response genes *addAB, comG*, and *comK*, respectively^30^. Each box in a promoter contains an adenine-thymine (AT) rich sequence (Fig. 1d). Such A-tracts are ubiquitous throughout all kingdoms of life, presumably due to their potential to curve DNA^31,32^. To assess the structural properties of the promoter, we engineered a set of *comG* promoters (18 bp spacer) that we site-specifically labelled with donor and acceptor fluorophores for confocal smFRET measurements (Extended Data Table 1). When we compared the experimental FRET efficiencies with theoretical values calculated for extended B-type DNA^29^, we noted substantial deviations across one box, suggesting an overall curved topology (Fig. 2a). Given that DNA curvature scales with DNA stiffness^33^, which is sensitive to the protonation state of the phosphate backbone, we repeated the experiment under more acidic conditions (pH 4.0, Fig. 2b). Indeed, we obtained reduced FRET values that better agreed with the calculated B-DNA profile. These results indicated that the *comG* promoter is curved at neutral pH. To probe ComK binding cooperativity, we labelled the *comG* promoter with a FRET-pair either at each box or within the spacer (Fig. 2c). In the absence of ComK, we find FRET values between 0.4 and 0.5. Upon addition of ComK, the FRET peaks shift to lower values for box 1 and box 2, equivalent to ∼3 Å distance increase, whereas the FRET efficiency increased for the spacer region, corresponding to a distance reduction of similar magnitude (Fig. 2c). Whereas the measured changes in FRET efficiencies were small, we were able to accurately derive relative populations of free and ComK-bound *comG* molecules by fitting our data to super- positions of two-state Gaussian peaks (Fig. 2c). We found that the fraction of ComK- *comG* complexes indeed increased in a sigmoidal fashion with ComK concentrations (Fig. 2d), in agreement with the cooperative mode of binding required by mathematical models of the phenotype switch^11,12,34^. Fits with the Hill equation (Fig. 2d)^35^ result in Hill exponents of *n* = 3.6 ± 0.1 for box 1 and *n* = 3.4 ± 0.3 for box 2. Independent controls with labelled ComK confirm that it is monomeric in solution (Extended Data Fig. 1-3). To interrogate coupling between box 1 and box 2, we created shorter single-box constructs (Fig. 2d, Extended Data Table 2). Remarkably, we obtained Hill coefficients of *n* = 1.7 ± 0.1 for box 1 and *n* = 1.5 ± 0.1 for box 2, which confirmed earlier reports of a 2:1 (protein:DNA) stoichiometry per box^30^. The twofold higher Hill exponents in the natural promoter show that the FRET-probes on one box also report on ComK-binding to the other box, which requires an allosteric communication between the boxes.

**Fig. 2.**
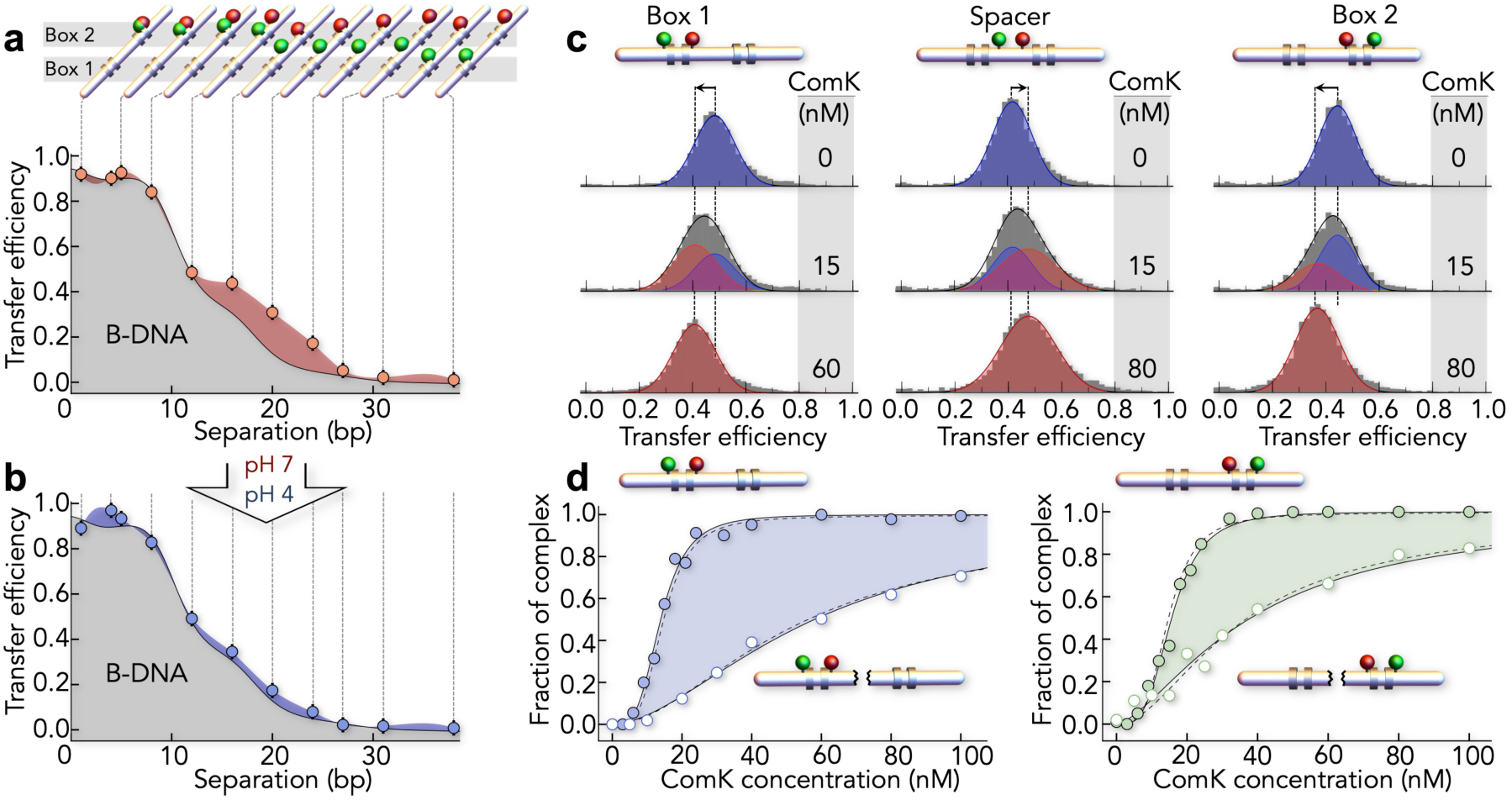
DNA-binding cooperativity of ComK probed with smFRET. (**a, b**) Mapping of distances in box 2 with FRET at higher (a, pH 7) and lower (b, pH 4) persistence length. Grey region indicates the expected dependence for B-DNA^29^. (**c**) SmFRET histograms of the *comG* promoter (93 bp) with an 18bp spacer indicate a change in DNA on ComK binding. FRET is probed in box 1 (left), in the spacer (middle), and in box 2 (right). Solid lines (black) are fits with super-positions of two Gaussian peaks for the free (blue) and ComK-bound (red) promoter. (**d**) Fraction of the ComK-DNA complexes probed in box 1 (left) and box 2 (right) in the presence (filled circles) and absence (empty circles) of the respective other box. The single-box constructs are 43 bp in length. Lines are fits with the Hill equation (solid) and with an induced-fit model (dashed).

## Structure of the complex

One classical mechanism to achieve allostery is DNA looping that allows contacts between proteins bound to different boxes^30,36^. Alternatively, ComK may bridge the boxes by interacting with the spacer, which could explain the distance decrease in the spacer upon binding of ComK (Fig. 2c, middle). To distinguish between these options, we used single-particle cryo-electron microscopy (cryo-EM) to overcome the strong aggregation tendency and low stability of ComK (Extended Data Fig. 4-6). We determined structures of ComK bound to promoters with spacers of 8 bp (*addAB:* 57 bp, 35.2 kDa) and 18 bp (*comG*: 93 bp, 57.5 kDa). Isolated DNA complexes of the expected size were readily identified in the cryo-EM micrographs (Extended Data Fig. 7) and 2D class averages revealed two major populations in both data sets (Fig. 3a, Extended Data Fig. 7). While the first population was pure promoter DNA, the second population contained DNA with extra density. Three observations are notable: (i) the extra density bound to DNA is confined to two locations that agree with the position of box 1 and 2, thus assigning this density to ComK; (ii) the DNA in these complexes is not looped, and (iii) only a few particles with ComK-density on only one box are found (Fig. 3b). The first observation demonstrates that ComK binds specifically to box 1 and 2 but not to the spacer between the boxes. Second, DNA-looping, as suggested in previous studies^30,37^, can be excluded as an allosteric mechanism by the 2D class averages. Indeed, we did not find an increase in FRET efficiency upon ComK binding to promoters labelled at the 5’ and 3’ end (Extended Data Fig. 10). Third, the lack of particles with ComK on only one box even under the vitrified conditions used in cryo-EM (Fig. 3b) suggests an all-or-none binding of the protein, in accord with the high cooperativity found with smFRET at room temperature (Fig. 2d). To determine the precise ComK-DNA stoichiometry, we reconstructed 3D-classes of the 18 bp spacer complex (Fig. 3c, Extended Data Fig. 7-9, 11). Although flexibility and preferred orientations limit the resolution to 8.9 Å, fitting of DNA into the 3D map enabled us to distinguish DNA and ComK (Fig. 3c). Comparison of the ComK mass in a box (42 ± 3 kDa) with that of a ComK monomer (22.6 kDa) yielded two ComK molecules per box (1.86 ± 0.13), i.e., four per promoter, in excellent accord with the Hill exponents (Fig. 2d). We further noted that ComK densities at individual boxes were continuous, which hints towards significant protein-protein contacts within a box (Fig. 3c). By contrast, we detected no ComK interactions across the spacer region, which suggests DNA-mediated allostery as source for the high binding cooperativity. To determine the allosteric mechanism, we analysed the DNA curvature of free (Fig. 3d) and bound promoters (Fig. 3e) and found a significant reduction in the presence of ComK, which hints at a potential mechanism. ComK is known to interact with the minor groove of DNA^30^, which is significantly narrower in A-tracts compared to B-DNA^38^. Indeed, a fit of our 3D density with an atomistic representation of the *comG* promoter suggests that ComK faces the minor groove by forming contacts with the thymine bases of the A-tracts (Fig. 3f, Extended Data Fig. 11). Altering the minor groove by ComK would lower DNA curvature around a box, thus affecting the width of the minor groove at the second box. Importantly, the model stipulates that DNA-mediated communication between boxes ought to be dependent on the spacer length.

**Fig. 3.**
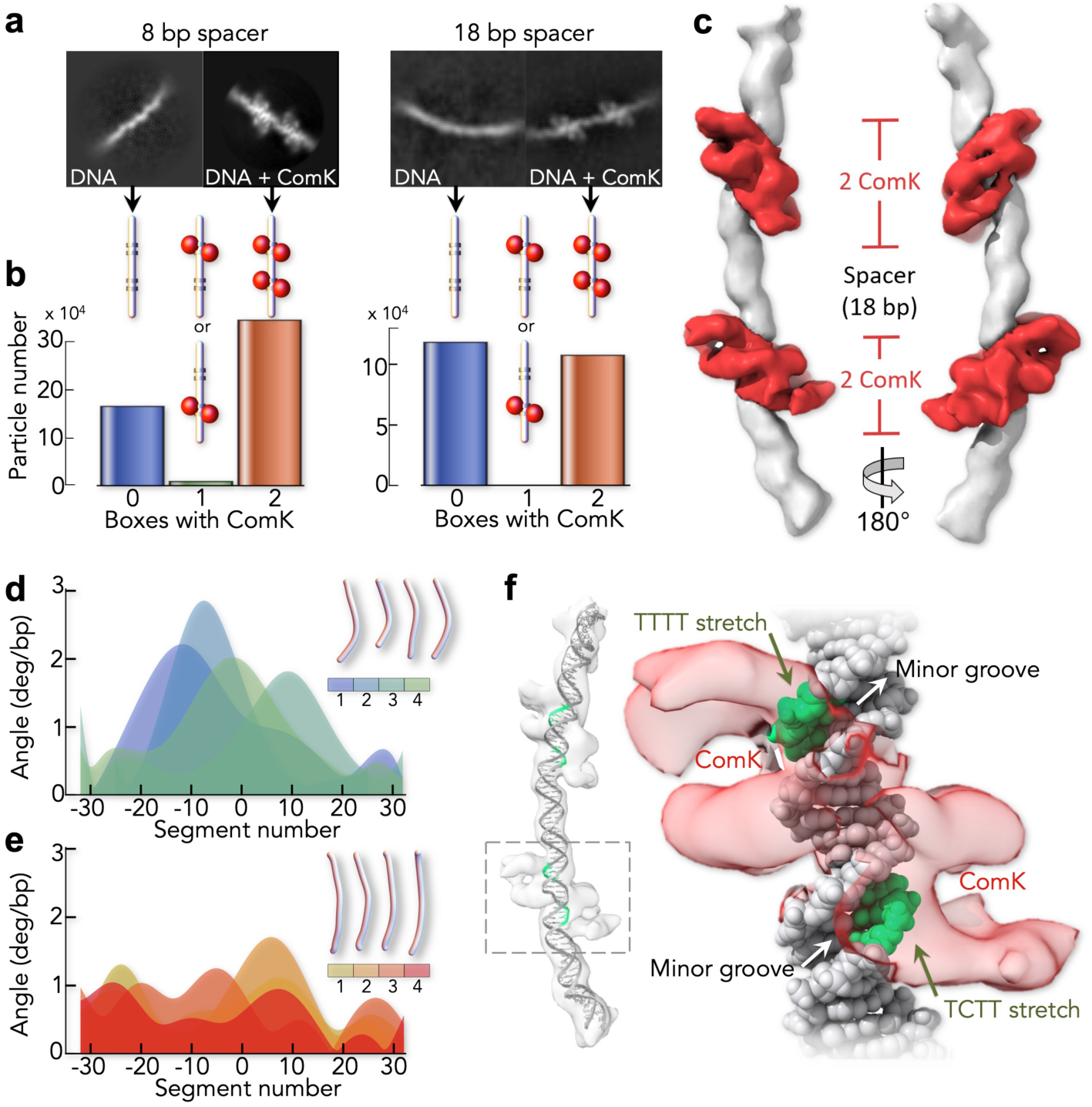
Single-particle cryo-EM analysis on ComK-DNA complexes. (**a**) Class averages of free DNA and the ComK-DNA complex for the promoters with an 8 bp (left) and 18 bp (right) spacer. (**b**) Distribution of particles with different boxes bound to ComK for the promoters with an 8 bp (left) and 18 bp (right) spacer. (**c**) 3D reconstruction of ComK bound to the promoter with an 18 bp spacer (Extended Data Fig. 7: 3D class 1 with 8.9 Å resolution). DNA (white) and ComK (red) densities were segmented following flexibly fitting of double-stranded DNA. The ComK volume at each box corresponds to ∼2 ComK molecules. (**d, e**) Curvature angles for free DNA (d) and the ComK-DNA complex (e) computed based on four 3D classes. Color code indicates the 3D classes with the overall shape of the DNA being indicated schematically. Angles between successive segments (segment length: 0.34 nm = 1bp) for different classes were aligned with respect to the middle of the DNA. Averaged over all segments, the difference in curvature angles is (0.3 ± 0.2)°/bp. (**f**) Flexible fitting of *comG* promoter DNA into 3D class 1 of the ComK-DNA complex (left) and orientations of ComK (red) relative to the A-tracts in a single box (right). ComK, shown at high iso-surface threshold, faces the thymine bases (green) in the minor groove (indicated). The DNA electron-density was subtracted and replaced by the fitted coordinates of the DNA model.

## Mechanical force transduction in DNA

To test this mechanism, we altered the spacer length and measured the effect on cooperativity using a binding model that is derived from the cryo-EM structures (Fig. 4a). Each box has two binding sites for ComK. The first ComK binds a box with the association constant *K*. The second molecule binds the same box with *σ*-fold increased affinity due to protein-protein contacts between the ComK molecules. Once the first box is saturated with ComK, the altered DNA curvature increases the affinity at the second box by a factor *J*. The quantity Δ*g*_*J*_ = - *k*_*B*_*T* log*J* is the DNA-mediated coupling free energy between two boxes. Using this model, we examined the coupling between boxes using the three promoters *addAB, comG*, and *comK* with spacer lengths of 8, 18, and 31 bp, respectively (Fig. 4b). When analysing the binding isotherms (Fig. 4b), we found a significant decrease in the Hill exponent with increasing spacer length along with a decrease of the coupling free energy (-Δ*g*_*J*_) from 5.8 ± 0.9 *k*_B_*T* (8 bp) to 1.9 ± 0.1 *k*_B_*T* (31 bp) (Fig. 4c). The result shows that spacer length is key for allostery, in line with the idea that reducing promoter curvature is the main origin for allosteric signal transmission. Compared to allosteric effects found previously in artificial promoters without curvature^14,23^, the coupling free energies are almost threefold larger, which converts to a fortyfold stronger effect on the association constants. Hence, curvature changes, though not mandatory^14,23^, greatly enhance DNA-mediated allostery.

**Fig. 4.**
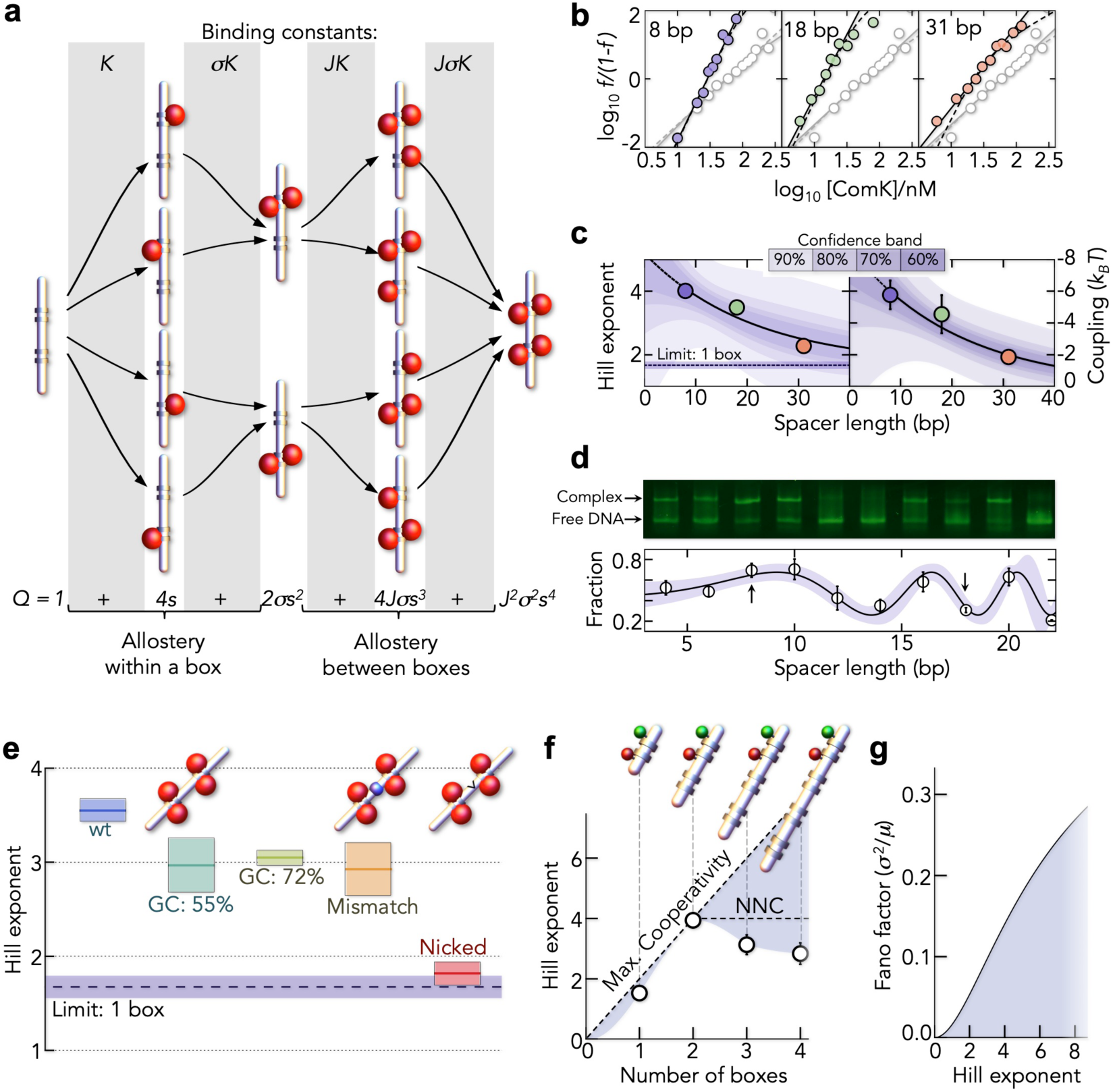
Tuning allostery. (**a**) Schematics of the binding model with DNA (grey) and ComK (red). A single box is first occupied with 2 ComK molecules before the second box is bound. The model contains a microscopic association constant *K* and the allosteric parameters *σ* and *J* due to allostery within a box and between boxes, respectively. Binding constants for each step are identical for all paths. (*Bottom*) Binding polynomial (*Q*) of the model with *s* = *K* [ComK]. (**b**) Hill-plots of the binding isotherms for promoters with varying spacer length (indicated) in comparison to an isolated box (grey). Solid lines are fits with the Hill-equation and dashed lines are fits with the model in A. (**c**) Hill exponents (*left*) and inter-box coupling free energies (*right*) as function of the spacer length. Solid lines are exponential fits and shaded areas indicate the confidence intervals of the fit. (**d**) Gel assay to screen ComK-binding to promoters with different spacer lengths with free DNA (lower band) and ComK-bound DNA (upper band) at a ComK concentration of 300 nM. (*Bottom*) Relative fraction of the ComK-DNA band as function of the spacer length. Arrows indicate promoters with 8 bp and 18 bp. Error bars indicate ±SD of independent triplicates. Solid line is a fit with a cosine function with spacer-dependent frequency. Shaded area is the 90% confidence band. (**e**) Hill exponents for *comG*-promoters with modified spacer. Colored lines are the mean and boxes indicate ±SD of at least two independent experiments. The dashed line indicates the limit of an isolated box. (**f**) Hill exponents of artificial promoters (8 bp spacer) with multiple boxes. The diagonal line indicates maximal cooperativity and the horizontal line indicates maximum nearest-neighbour cooperativity (NNC). Error bars represent the error of the fit. (**g**) Increase in the noise of the fraction of ComK-bound promoters with increasing cooperativity. ComK copy numbers were sampled (*n* = 10^5^) from a Poisson distribution with a mean of 9 molecules per cell (∼15 nM). Variance (*σ*^2^) and mean (*μ*) of *f* (Fig. 1c) were computed for different Hill exponents with *K*_*Hill*_ = 15 nM.

A theory of DNA bending predicts an exponential decay of Δ*g*_*J*_ with increasing length of the spacer^18^. The decay length (ξ) is given by *ξ* = (*k*_B_*T l*_*p*_/*f*)^1/2^ with the persistence length *l*_*p*_ = 40 nm^39^ and a tension force *f*. We found *ξ* = 21 ± 5 bp and *f* = 0.15 ± 0.07 pN/bp (Fig. 4c). Given the known energetic costs of DNA bending^40^, we estimated a 0.2 ± 0.1°/bp change in curvature, which agrees well with the change determined from the cryo-EM structures (Fig. 3d,e). Yet, the coarse-grained model does not account for the helical periodicity of DNA. Indeed, a gel assay shows that the fraction of ComK-DNA complexes oscillates with the spacer length between the boxes (Fig. 4d) and it is likely that also the cooperativity will depend on whether two boxes are in-phase or out-of-phase with respect to the helical DNA-turns.

## Tuning allostery via DNA sequences

To understand the sequence determinants of allostery, we altered the spacer sequence in *comG* by increasing its GC-content, thus increasing DNA stability. The GC-content of the native spacer (40%) is close to the genome average^41^. We generated two random spacer sequences of higher GC content (55% and 72%, Extended Data Table 2). Indeed, increasing GC-content lowered cooperativity (Fig. 4e). Nevertheless, the Hill exponents remained above the values found for the isolated boxes, suggesting that spacer sequences fine-tune allostery at best. Next, we tested whether allostery requires a correct base pairing by introducing a mismatch of four bases at the centre of the spacer. However, even though the Hill exponent is reduced compared to wild-type *comG*, it is comparable to those found for increased GC contents (Fig. 4e). Remarkably, this result suggests that the allosteric communication between boxes does not stringently require a continuous base pair stacking in the spacer, thus pointing at the DNA backbone as the major determinant for allostery. Indeed, when we introduce a nick in the spacer, cooperativity is abolished and we obtain a Hill exponent of *n* = 1.8 ± 0.1, i.e., close to the value found for the isolated boxes (Fig. 4e). We therefore conclude that curvature-induced tension between the boxes propagates mainly via an intact DNA backbone but it can be fine-tuned by the spacer sequence and length.

Finally, we tested promoters with multiple boxes. In fact, the *comG* promoter contains an A-tract located 10 bp upstream of the first box (Extended Data Table 2)^37^. Although this region is not occupied in our cryo-EM structure due to its imperfect sequence, increasing the number of perfect boxes may be a simple strategy to further boost cooperativity. However, even the presence of 4 boxes with short spacing (8 bp) does not increase the Hill exponent beyond the value found for the promoter with two boxes. This implies that curvature changes do not propagate across multiple boxes but rather mediate nearest-neighbour communication (Fig. 4f).

## Conclusions

Our results show that DNA-mediated allostery generates high cooperativity in transcription factor binding via mechanical deformations of the DNA across distances of 6 nm (18 bp). Notably, this mechanism is a built-in feature of natural ComK promoters to filter copy number fluctuations of ComK (Fig. 1b,c): copy numbers below the midpoint of the Hill curve leave the promoter unoccupied, those above the midpoint cause near full occupation. With increasing cooperativity, this filter increases the variability (noise) of free and bound promoters (Fig. 4g), which would also affect the ratio of vegetative and competent cells in a population. However, even if genetic circuits function in the deterministic regime of high copy numbers, sigmoidal dose responses of gene expression activity with transcription factor concentrations are key for their dynamics^42-44^. Altering the steepness of this response, i.e., the molecular cooperativity, will unambiguously alter these dynamics^10,45^. Hence, besides the architecture of wiring genes^46,47^, designing dose-response curves via promoter sequences will provide a new level of engineering the dynamics of gene networks in the future.

## Data and materials availability

Cryo-EM maps and atomic coordinates have been deposited in the Electron Microscopy Data Bank (EMDB) and Protein Data Bank (PDB) under accession codes EMD-11022 and 6Z0S, respectively.

## Author Contributions

G.R. and H.H. designed research. G.R. and F.W. labelled DNA constructs. G.R. and H.H. performed and analysed single-molecule experiments. G.R. and N.E. performed cryo-EM work and analysed these data. H.R. conducted the fitting of the 3D map with an atomic model of the DNA. H.H. wrote the paper with the help of all authors.

## Acknowledgements

We enjoyed the critical discussions and helpful comments of many colleagues. Our thanks go to Deborah Fass, Gilad Haran, Amnon Horovitz, Benjamin Schuler, and Philipp Selenko. We also thank Harry Greenblatt for computational support as well as Christian Dubiella and Ronen Gabizon for their help with the LC-MS. This research was supported by the Israel Science Foundation (grant no. 1549/15), the Benoziyo Fund for the Advancement of Science, the Carolito Foundation, the Gurwin Family Fund for Scientific Research, the Leir Charitable Foundation, and the Koshland family. The work was further supported by the Irving and Cherna Moskowitz Center for Nano and Bio-Nano Imaging at the Weizmann Institute of Science.

## Notes

### Competing Interest Statement

The authors have declared no competing interest.

