## Supplementary Information for "Allostery through DNA drives phenotype switching"

### METHODS

**DNA labelling and duplex preparation.** Single-stranded DNA constructs (Extended Data Table 1) were synthetized and HPLC-purified by the Keck Biotechnology Resource Laboratory (Yale School of Medicine). For labelling with the amino-group reactive dyes AlexaFluor 488 (donor) and AlexaFluor 594 (acceptor) NHS ester (ThermoFisher), thymine or adenine bases at the labelling positions contained an amino modifier C6-dT/dA CE phosphoramidite (Glen Research) (Extended Data Table 1-2). In constructs used for smFRET ComK-binding experiments, the positions of the modified A- and T-bases were chosen such that nucleotides were avoided that had previously been shown to be protected by ComK-binding[1]. For labelling, we followed the procedure by Masoud *et al.*[2] with the exception that the labelling reaction was incubated for 1 hour at 45°C. The donor and acceptor dyes were used to label the forward and reverse ssDNA strands, respectively. Reversed phase HPLC was used to remove unreacted dye, and unlabelled ssDNA. To this end, a ZORBAX Eclipse Plus C18 (3.5 μm) column (Agilent) was used, equilibrated with TEAA buffer (0.1M trimethylamine buffered with acetic acid glacial to pH 6.5 and supplemented with 5% acetonitrile). A gradient of 7% - 30% acetonitrile over 40 ml was used for the separation.

Labelled fractions were dried overnight (SpeedVac) and dissolved in 50 μl $ddH_2O$. DNA and dye concentrations were quantified by UV-VIS photometry using the extinction coefficients of the free dyes $\varepsilon_{495}$=73,000 and $\varepsilon_{590}$=92,000 $M^{-1}$ $cm^{-1}$, respectively. Only fractions containing 1:1 ratio of dye:ssDNA were used for smFRET experiments. Annealing of ssDNA donor and acceptor strands for smFRET experiments was obtained by mixing 2 pmol donor-labelled ssDNA with 3 pmol acceptor-labelled ssDNA in 50 mM Tris pH 7.5, 0.1 M NaCl, 3 mM $Mg_2Cl$. The samples were heated to 95°C and cooled gradually over 1 h in a PCR cycler. Unlabelled DNA duplexes for cryo-EM and EMSAs (electrophoretic mobility assays) were obtained by mixing a 1:1 ratio of forward and reverse ssDNA strands using the same annealing procedure.

**Expression and purification of ComK.** The sequence of ComK (Uniprot ID: P40396) codon-optimized for expression in *E. coli* (Genscript) (Extended Data Table 3) was inserted into the NcoI and HindIII sites of a pET28b vector, and the protein was expressed in inclusion bodies (IBs) in BL21(DE3). Bacteria were grown in 4 l LB medium containing 50 μg/ml Kanamycin. Cultures were induced at $OD_{600}$ 0.7 with 0.3 mM IPTG. After 4h at 37°C, the cells were harvested by centrifugation and the cell pellets were re-suspended in lysis buffer (50 mM Tris pH 8.0, 10 mM EDTA, 100 μM PMSF, 1.2 μg/ml leupeptin and 1 μM pepstatin A). After sonication (70% power, pulse on 2s, pulse off 10 s, 60 cycles, on ice) and centrifugation (48,200 g, 30 min at 4°C), the IB precipitate was mixed with 15 ml of 60 mM EDTA, 6% Triton, 1.5 M NaCl, homogenized and incubated at 4°C for 1 hour to solubilize membrane debris. After centrifugation (48,200 g, 30 min at 4°C), the resulting IB precipitate was supplemented with 0.1 M Tris pH 8, 1 mM EDTA, homogenized and centrifuged using the same conditions. Finally, the IB-pellet was re-solubilized (>2h, 4°C) in 50 mM Tris pH 8.0, 6 M GdmCl, 10mM imidazole, containing 10mM DTT. After complete resolubilization, the buffer was exchanged to 50 mM Tris pH 8.0, 6 M GdmCl, 10mM imidazole using a HiPrep desalting column (26/10, GE) to remove DTT. The protein containing fractions were then loaded on a HisTrap Ni-NTA column (5 ml, GE) and the protein was eluted with 50 mM Tris pH 8.0, 6 M GdmCl, 0.5 M imidazole using a linear gradient of 3 ml/min for 10 min. The eluted protein (20 ml) was supplemented with 0.1mg/ml HRV3C and 5 mM DTT and the solution was dialyzed overnight (4°C) against 300 volumes of cleavage buffer (50 mM Tris and 100 mM L-Arg at pH 7, 1mM DTT). HRV3C protease refolds efficiently under these conditions. Due to the strong aggregation propensity of ComK, the protein partially precipitates during this procedure. The protein was fully precipitated by the addition of an equivalent volume of saturated ammonium sulfate solution. The precipitate was collected by centrifugation (20,000 rpm, for 20 min at 4°C) and the protein pellet was dissolved in 50 mM Tris pH 8.0, 6 M GdmCl, 10mM Imidazole. The cleaved His-tag and HRV3C protease were removed using Ni-NTA affinity chromatography (5 ml HisTrap column, GE). Cleaved, i.e., untagged protein was obtained in the flow-through fractions while the purification tags and HRV3C protease were retained on the column. Finally, the buffer of the ComK containing fractions was exchanged to 0.5 M L-Arg pH 7.3 using a HiTrap desalting column (5 ml, GE). The protein concentration was determined using an extinction coefficient of $\varepsilon_{280}$ = 24,660 $M^{-1}$ $cm^{-1}$ computed from the amino acid sequence of ComK. Since ComK contains 4 cysteines, we added 10 mM DTT and stored the samples at -80ºC. For mass determination, the purified ComK in 0.5 M L-Arg was injected into a combined LC-MS system (Waters ACUITY UPLC class H) equipped with a C4 column (300 Å, 1.7 µm, 21 mmx100 mm) and the mass was determined in positive ion mode using electrospray ionization using a desolvation temperature of 500 °C with flow rate of 1000 liter/hour. The voltage was 0.69 kV for the capillary and 46 V for the cone. Spectra, de-convoluted with the MaxEnt1 software, resulted in a single peak with a mass of 22,652±10 Da, in good agreement with a calculated mass of 22,640 Da.

**Labelling of ComK for 2fFCS experiments**. Singly labelled ComK is required to determine the oligomeric state of ComK in solution using 2fFCS[3]. Since ComK contains four intrinsic cysteine residues at positions 31, 39, 55, and 102 (Extended Data Table 3), our approach was to replace them by alanine. However, the resulting $\Delta$Cys variant was incapable of binding DNA at protein concentrations ≥ 100 nM as demonstrated in smFRET experiments using doubly labelled *comG* (Extended Data Fig. 1). The result surprises given the facts (i) that ComK is a cytosolic protein that acts in a reducing environment and (ii) that all our smFRET experiments with wildtype ComK are performed under strongly reducing conditions (20 mM DTT). The result shows that the reduced cysteine residues are either crucial for the structural integrity of ComK, e.g., as metal binding centres, and/or are key for the interaction with the DNA.

To obtain active and fluorescently labelled ComK for 2fFCS measurements, we therefore introduced an additional well-accessible cysteine at the N-terminus of ComK for labelling. After purification and HRV3C-cleavage, we added 1.4 mol equivalents of AlexaFluor488 C5 maleimide to 46 µM ComK in 0.5 M Arginine-HCl, pH 7.3. After 30 min at room temperature, the reaction was quenched by the addition of 10 mM DTT and the buffer was exchanged to 50mM Tris-HCl pH 8.0, 6M GdmCl. Unlabelled ComK was separated from labelled ComK using reversed-phase chromatography (RP-C18 column) in 0.1% TFA in $ddH_2O$. The protein was eluted using a gradient of 25-50% acetonitrile at a flow rate of 1 ml/min for 40min. The fraction containing labelled ComK was first lyophilized and then dissolved in 50mM Tris-HCl pH 8.0, 6M GdmCl buffer. A UV-Vis quantification confirmed a 1:1 ratio of ComK to dye.

**Optimizing solution conditions to prevent ComK aggregation**. Our results show that ComK aggregates at concentrations as low as 2 μM (Extended Data Fig. 5). High concentrations of L-Arg (0.5 M) effectively prevent this aggregation up to a concentration of >100 μM. However, our experiments also show that such high L-Arg concentrations suppress the binding of ComK to DNA. A quantification of the aggregation of ComK using UV/VIS photometry revealed that at least 50 mM L-Arg (Extended Data Fig. 5) is required to ensure soluble ComK up to a concentration of 2 μM. Even though the L-Arg concentration exceeds the cytosol concentration of 0.57 mM (*E. coli*)[4], our control experiments show that the conformation of the DNA (*comG*) is unaffected by L-Arg up to a concentration of 100 mM, and ComK is fully active in binding DNA (Extended Data Fig. 5). We therefore used 50mM L-Arg in all smFRET and cryo-EM experiments. Notably, L-Arg is known to stabilize proteins, thus preventing aggregation[5], however, it does not solubilize pre-formed aggregates in our hands.

**SmFRET and 2fFCS experiments**. All smFRET and 2fFCS experiments with ComK were performed at 20 mM Tris pH 7.0, 50 mM L-Arg, 5 mM $MgCl_2$, 150 mM KCl, 20 mM DTT, 0.001% Tween-20. In contrast, the smFRET experiments in which we map the curvature of the comG promoter (Fig. 2b,c) were performed in 20 mM sodium phosphate pH 7 or 4, 100 mM β-mercapto-ethanol, 0.001% Tween-20, which allowed an easier adjustment of the solution pH. The ionic strength was kept constant at 100 mM in these experiments by the addition of NaCl. The concentration of labelled DNA was 30 pM.

All smFRET experiments were performed with a MicroTime 200 confocal microscope (PicoQuant) equipped with an Olympus IX73 inverted microscope. Briefly, we used linearly polarized light from a 485 nm diode laser (LDH-D-C-485, PicoQuant) and unpolarized light at 594 nm from a supercontinuum light source (Solea, PicoQuant) to excite donor and acceptor alternatingly with a total repetition rate of 40 MHz. This pulsed-interleaved excitation (PIE) scheme[6] was used to distinguish molecules with only donor dye from those carrying both dyes but with a large inter-dye distance. The excitation beam was guided through a major dichroic mirror (ZT 470-491/594 rpc, Chroma) to a 60x, 1.2NA water objective (Olympus) that focuses the beam into the sample. The sample was placed in a home-made cuvette with a volume of 50 µl made of round quartz cover slips of 25 mm diameter (Esco Optics) and borosilicate glass 6 mm diameter cloning cylinder (Hilgenberg) using Norland 61 optical adhesive (Thorlabs). All measurements were performed at a laser power of 100 µW (485 nm) and 20 µW (594 nm) measured at the back aperture of the objective. Photons emitted from the sample were collected by the same objective and after passing the major dichroic mirror (ZT 470-491/594 rpc, Chroma), the residual excitation light was filtered by a long-pass filter (BLP01-488R, Semrock) and focused on a 100 µm pinhole. The sample fluorescence was detected with two channels. Donor and acceptor fluorescence was separated via a dichroic mirror (T585 LPXR, Chroma) and each color was focused onto a single-photon avalanche diode (SPAD) (Excelitas) with additional bandpass filters: FF03-525/50, (Semrock) for the donor SPAD and FF02-650/100 (Semrock) for the acceptor SPAD. The arrival time of every detected photon was recorded with a HydraHarp 400M time-correlated single photon counting module (PicoQuant) and stored with a resolution of 32 ps.

For 2fFCS (two-focus fluorescence correlation spectoscopy)[3], the sample was excited with two orthogonally polarized lasers at 485 nm (LDH-D-C-485, PicoQuant). The beams of both lasers were combined and laterally shifted by a Nomarsky prism at the back aperture of the objective. Emitted light passed through a 150 µm pinhole. Afterwards, the emitted light was split using a polarizing beam splitter and focused on two separate SPAD detectors. The distance between the two resulting foci was experimentally determined using the known Stokes radii ($R_S$) of the calibration samples: the dye Oregon Green ($R_S = 0.6$ nm, 0.36 kDa) and the proteins Aprotinin ($R_S = 1.29$ nm, 6.5 kDa), RNaseA ($R_S = 1.73$ nm, 13.7 kDa), Carboxy Anhydrase ($R_S = 2.19$ nm, 29 kDa), Ovalbumin ($R_S = 2.65$ nm, 44 kDa), Conalbumin ($R_S = 3.1$ nm, 75 kDa) (Extended Data Fig. 1). For calibration, all proteins were labelled with AlexaFluor488 NHS ester at primary amino groups of lysine residues. The autocorrelation functions of the signal of each focus and the two cross-correlation functions were fit globally with a model that contained one diffusion component and two exponential decay components to account for the triplet dynamics of the dyes (Extended Data Fig. 2a). The global fits were performed iteratively to minimize the squared difference between the calibration values and the values determined with 2f-FCS. Subsequently, the Stokes radius of labelled ComK was determined in the absence (Extended Data Fig. 2b) and presence (Extended Data Fig. 3) of increasing amounts of unlabelled ComK. All measurements were performed at 23°C.

**Determination of the thermodynamic stability of ComK**. To understand the strong aggregation tendency of ComK, we used 2fFCS to determine the stability of ComK under physiological conditions. To this end, we denatured ComK at different concentration of the denaturant GdmCl (guanidinium chloride). With increasing concentrations of GdmCl, the Stokes radius increases in a cooperative manner (Extended Data Fig. 4). To obtain the stability, we fit the data using a two-state model of protein folding. Assuming a spherical shape of folded and unfolded ComK, the average Stokes radius $\langle R \rangle$ at any concentration of GdmCl is given by

$$\langle R \rangle = \left\{ \frac{3}{4\pi} \left[ f v_f + \left( 1\text{-}f \right) v_u \right] \right\}^{1/3} \quad (1)$$

Here, $f$ is the fraction of folded molecules, $v_f$ and $v_u$ are the volumes of folded and unfolded molecules, respectively. Since it is known that the dimension of unfolded polypeptide chains responds sensitively to the concentration of denaturants[7-9], we assume that the volume of the unfolded state changes linearly with the concentration of GdmCl ($x$) according to

$$v_u = m_u + n_u \quad (2)$$

with the slope $m_u$ and the intercept $n_u$. The fraction $f$ for a two-state system is given by

$$f = \frac{e^{-\Delta G/RT}}{1 + e^{-\Delta G/RT}} . \quad (3)$$

Here, $R$ is the ideal gas constant, $T$ is the temperature in Kelvin, and $\Delta G$ is the free energy difference between folded and unfolded molecules. We further assume a linear free energy relationship of the type

$$\Delta G = mx + \Delta G_0 . \quad (4)$$

The data in Extended Data Fig. 4 were fit using a combination of eq. 1-4. The fit results in free energy of stabilization of $\Delta G_0$ = -8.8 ± 0.3 kJ/mol.

**SmFRET data analysis.** Fluorescence bursts from individual molecules were determined by combining successive photons separated by inter-photon times of < 100 µs into one burst with $n_D$ and $n_A$ photons counted in the donor and acceptor detection channels, respectively. Identified bursts ($n_A + n_D > 80$) were corrected for background, differences in quantum yields of donor and acceptor, different collection efficiencies in the detection channels, cross-talk, and direct acceptor excitation as described previously[10], thus giving the corrected photon counts $n_A'$ and $n_D'$. Transfer efficiency histograms were computed according to

$$E = \frac{n_A'}{n_A' + n_D'} \quad (5)$$

In addition, only molecules containing active acceptor and donor dyes were included in the analysis. To this end, we computed the donor-acceptor stoichiometry ($S$) for each burst according to

$$S = \frac{n'_{DD} + n'_{DA}}{n'_{DD} + n'_{DA} + \gamma\left(n'_{AD} + n'_{AA}\right)} \quad . \qquad (6)$$

Here, $\gamma$ is a correction factor to account for the different excitation intensities for donor and acceptor. Furthermore, the first subscript indicates the emission and the second subscript indicates the excitation. Only molecules with $S < 0.8$ were used for constructing smFRET histograms. FRET histograms were fitted with a combination of two Gaussian distributions[11]. The width and position of the FRET-peaks were fixed to minimize the number of free fitting parameters and the area under the histogram curve for each subpopulation was determined using numerical integration to obtain the fraction of ComK-bound promoter. For distance calculations based on mean transfer efficiencies, the Förster radius measured in water is $R_0 = 5.4$ nm [7].

**Determination of the fraction of ComK-bound promoters from smFRET.** To determine the fraction of ComK-bound promoters, we fit the FRET-histograms with a superposition of Gaussian or log-normal distributions[11]. To reduce the number of fitting parameters, we first fit the FRET histogram in the absence of ComK and in the presence of saturating concentrations of ComK with a single peak. The position, width, and, in the case of log-normal distributions, also asymmetry, are then fixed to these values. All FRET-histograms at intermediate ComK concentrations are then described by a linear superposition of the peaks determined at the two limiting conditions. The fraction of ComK-bound promoter is finally obtained by the areas of the two peaks (free and bound promoter).

**Calculation of the FRET values for B-DNA.** To determine the transfer efficiency profile of B-DNA (Fig. 2b,c), we followed the work of Clegg *et al.*[12] and Wozniak *et al.*[13]. Keeping the nomenclature of Wozniak *et al.*, the mean distances $R_{DA}$ of donor ($D$) – acceptor ($A$) pairs on B-DNA is given by

$$R_{DA} = \sqrt{\left(L + \Delta_{bp} z_{bp}\right)^2 + r_A^2 + r_D^2 - 2 r_A r_D \cos\left(\alpha + \Delta_{bp} \beta_{bp}\right)} \qquad (7)$$

Here, $\alpha$ is the offset angle between the fluorophores if they were both bound to the same base. $L$ accounts for the fact that projections of the centers of $D$ and $A$ onto the helix axis do not necessarily coincide with the position of bases. The values $r_A$ and $r_D$ are the distances from the helical axis due to the dye linkers, $z_{bp}$ is the increase along the axis per base pair, and $\beta_{bp}$ is the increase in D-A angle per base pair. For dyes with a similar size of those used here, Wozniak *et al.* found $L = 0.617$ nm, $z_{bp} = 0.338$ nm, $r_A = 1.177$ nm, $r_D = 1.247$ nm, $\beta_{bp} = 36°$, and $\alpha = 89.8°$ for B-DNA and we used the same parameters to compute the transfer efficiency via

$$E_{mp}=\frac{R_0^6}{R_0^6+R_{DA}^6} \quad (8)$$

Since our experimentally measured transfer efficiencies are an average over the positional distribution of the dyes due to the flexibility of their C5-linkers, the measured transfer efficiencies $\langle E\rangle$ were first converted according to the values expected for the mean positions according to[13]

$$E_{mp}=0.008+0.679\langle E\rangle+1.470\langle E\rangle^2-1.141\langle E\rangle^3. \quad (9)$$

**Thermodynamic ComK-DNA binding model.** To determine coupling free energies between the binding boxes of ComK, we fit the data using a mechanistic binding model (Fig. 4a). In this model, we denote the promoter as $P$ and the four binding sites are denoted by subscripts $P_1 \dots P_4$. Here, the first two binding sites are in box 1 and the second two binding sites are in box 2. We further denote ComK as $X$ such that the concentration of the promoter with one ComK bound to the first site is $[P_1X]$. The concentration of free ComK is $x$. The initial binding of a ComK to each site is given by four association reactions with identical association constant $K$ defined by

$$K=\frac{[P_1X]}{[P]x}=\frac{[P_2X]}{[P]x}=\frac{[P_3X]}{[P]x}=\frac{[P_4X]}{[P]x} \quad (10)$$

The second ComK binds exclusively to a box that already contains a bound ComK. We introduced this constraint based on the determined cryo-EM structure that indicates substantial protein-protein contacts between ComK molecules in one box. Hence, binding the second ComK gives the relations

$$\sigma K=\frac{[P_{12}X_2]}{[P_1X]x}=\frac{[P_{12}X_2]}{[P_2X]x}=\frac{[P_{34}X_2]}{[P_3X]x}=\frac{[P_{34}X_2]}{[P_4X]x} \quad (11)$$

The factor $\sigma$ accounts for the higher affinity of binding a second ComK to a box due to allostery via protein-protein contacts. The third ComK now binds to a different box, thus leading to

$$JK=\frac{[P_{134}X_3]}{[P_{34}X_2]x}=\frac{[P_{234}X_3]}{[P_{34}X_2]x}=\frac{[P_{123}X_3]}{[P_{12}X_2]x}=\frac{[P_{124}X_3]}{[P_{12}X_2]x} \quad (12)$$

where the factor $J$ accounts for the inter-box cooperativity. Finally, the last ComK molecule binds to the remaining site that now includes intra-(σ) and inter-($J$) box cooperativity

$$J\sigma K=\frac{[P_{1234}X_4]}{[P_{234}X_3]x}=\frac{[P_{1234}X_4]}{[P_{134}X_3]x}=\frac{[P_{1234}X_4]}{[P_{124}X_3]x}=\frac{[P_{1234}X_4]}{[P_{132}X_3]x} \qquad (13)$$

Replacing $s=Kx$ for clarity and with $[P_0]$ as the total concentration of promoter DNA, we obtain the mass conservation equation

$$[P_0]=[P]\left(1+4s+2\sigma s^2+4J\sigma s^3+J\sigma^2 s^4\right), \qquad (14)$$

which gives the binding polynomial shown in figure 4a

$$Q=1+4s+2\sigma s^2+4J\sigma s^3+J\sigma^2 s^4. \qquad (15)$$

The fraction of fully bound promoter is therefore given by

$$f=\frac{J^2\sigma^2 s^4}{1+4s+2\sigma s^2+4J\sigma s^3+J^2\sigma^2 s^4}\,. \qquad (16)$$

With the same logic, we obtain for the isolated boxes

$$f=\frac{\sigma s^2}{1+2s+\sigma s^2}\,. \qquad (17)$$

Notably, these equations only hold under conditions at which the concentration of DNA is very low compared to $K^{-1}$, which, with $[P_0]$=30pM is given at all our experimental conditions. To fit our binding data, we first fit the data of the isolated boxes 1 and 2 globally using eq. 17 (Fig. 2d). Here, the value $\sigma=e^{-\Delta g_\sigma}$ was a global parameter whereas $K=e^{-\Delta g_K}$ was a local parameter that differed between box 1 and box 2. These fits provide the free energy changes $\Delta g_\sigma$ and $\Delta g_K$ in (Extended Data Table 4). To reduce the number of parameters in eq. 9, we kept the value of $\Delta g_\sigma$ in all fits of promoters with two boxes and expressed the coupling between two boxes as $J=e^{-\Delta g_J}$ with the coupling free energy $\Delta g_J$ (Extended Data Table 4). For promoters in which we mapped both boxes (*ab1* and *comG*), we fit the data using a global value for $\Delta g_J$ (Fig. 2e and 4b). To allow an easier comparison of the cooperativity in the different constructs, all data were also fitted using the standard Hill-equation

$$f=\frac{x^n}{K_{Hill}^n+x^n} \qquad (18)$$

where $n$ is the Hill exponent that is an empirical measure for cooperativity, and $K_{Hill}$ is the effective dissociation constant.

**Gel retardation assay.** The gel retardation assay, also termed electrophoretic mobility shift assay (EMSA), is capable of providing a qualitative estimate of ComK–DNA binding. The forward strand (Extended Data Table 5) and the reverse strand, at a final concentration of 10 μM were mixed in the following buffer: 50 mM Tris pH 7.5, 0.1 M NaCl, 3 mM MgCl. The mixture was heated to 95ºC and slowly cooled in a PCR cycler as previously described. The reaction mixture contained 25 nM dsDNA, 300 nM ComK, in the following buffer: 20 mM Tris pH 7.0, 50 mM L-Arg, 5 mM $MgCl_2$, 150 mM KCl, 10 mM DTT, 0.001% Tween-20. The reaction was incubated for 20 min and supplemented with glycerol to a final concentration of 10%. The reaction was loaded on a 5% native PAGE, composed of 4.2 ml of 30% polyacrylamide 29:1, 8.3 ml of 1.5 M Tris-HCl pH 8.8, 12.2 ml $ddH_2O$, 150 μl of 10% Ammonium persulfate and 150 μl TEMED. The gels were ran in a Hoefer system at 100 V for 30 min, under TAE running buffer. The gels were than incubated in 100 ml $ddH_2O$ and a single drop of 0.625 mg/ml Ethidium Bromide, for 10 min. The gels were then washed and imaged with a Typhoon FLA 9500 scanner (GE).

**Sample preparation for cryo-EM.** ComK at a concentration of 40 μM (500 μl) in 50 mM Tris pH 8.0, 6 M GdmCl, 10mM Imidazole was injected on a HiTrap desalting column (5 ml, GE Healthcare) pre-equilibrated with 0.5 M Arginine-HCl, pH 7.3. The protein containing fractions (32 μM) were then supplemented with 10 mM DTT to prevent the oxidation of the intrinsic cysteine residues of ComK. The ComK-DNA complexes were formed by quickly mixing 2.5 μl of ComK (32 μM) and 0.6 μl of DNA (20 μM *addAB* or *comG*) with 22 μl 20 mM Tris pH 7.0, 5 mM $MgCl_2$, 150 mM KCl, 3mM DTT, 0.001% Tween-20. Immediately, after mixing, 3.5 μl of this mixture was transferred to C-flat 2/2 200 mesh holey carbon grids (Electron Microscopy Sciences). For addAB, 2.5 μl were transferred to Quantifoil 0.6/1 200 mesh holey carbon grids. The grids for both samples were glow discharged before applying the sample, blotted for 2.5 s at 4°C and 100% humidity, and plunge frozen in liquid ethane cooled by liquid nitrogen using a Vitrobot plunger (Thermo Fisher Scientific).

**Cryo-EM data acquisition.** Cryo-EM data sets were collected on a Titan Krios G3i transmission electron microscope (Thermo Fisher Scientific) operated at 300 kV. The ComK-*comG* data set comprised 5210 movies recorded on a Falcon 3EC direct detector in counting mode at a nominal magnification of 75,000×, which corresponds to a physical pixel size of 1.09 Å. The dose rate was set to 0.84 $e^-$/pixel/s and the total exposure time was 42.7 s, which resulted in an accumulated dose of 30.2 $e^-/Å^2$. Each Movie was fractionated into 30 frames. The nominal defocus range was -0.6 to -2.2 μm. Imaging was done using an automated low dose procedure implemented

in EPU software (Thermo Fisher Scientific) in which stage navigation was used to navigate to hole centers and image shift was used to target 4 distinct imaging locations within each hole.
The ComK-*addAB* data set comprised 12,338 movies recorded on a K3 direct detector (Gatan) at the end of BioQuantum energy filter (Gatan) using a slit of 20 eV. Movies were recorded in counting mode at a nominal magnification of 105,000×, which corresponds to a physical pixel size of 0.82 Å. The dose rate was set to 18.2 $e^-$/pixel/s and the total exposure time was 3.25 s, resulting in an accumulated dose of 88 $e^-/Å^2$. Each movie was fractionated into 75 frames of 0.044 s. The nominal defocus range was -1 to -1.8 μm. SerialEM[14] was used for automated low dose data collection.

**Single particle cryo-EM image processing.** Image processing was performed using CryoSPARC software[15]. Movie frames were aligned using patch motion correction, followed CTF estimation using CTFFIND4 [16] module within CryoSPARC. In total, 3909 ComK-*comG* images, which featured resolution better than 8.0 Å and cross-correlation fit score higher than 0.05 were selected for further processing. The processing procedure is outlined in Extended Data Fig. 7. A subset of ~2000 particles were manually selected and subjected to a reference-free 2D classification followed by automated picking using the newly generated 2D class averages. About 600,000 automatically picked particles were extracted, binned 6x6 (60 pixel box size, 6.54 Å/pixel), and subjected to multiple rounds of reference-free 2D classification in order to clean the data set. It is difficult to distinguish the ComK-bound complexes from the free DNA in the raw images, however they are easily distinguished after 2D classification (Extended Data. 7). Thus, the 2D class averages from the mixed data set were separated into two sets of reference, ComK-DNA and free DNA. The selected micrographs were then subjected to automated picking, each time using one of the reference sets, resulting in two separate data sets, ComK-DNA (549,315 particles) and free DNA (444,641 particles). The two data sets were cleaned by multiple rounds of 2D classification, leaving 135,116 particles and 156,479 particles in the ComK-DNA and free DNA data sets respectively. 2D classification was followed by *ab-initio* 3D reconstructions, first into a single class, and then to multiple classes (60 pixel box size, 6.54 Å/pixel). We tried different numbers of 3D classes that may account for the heterogeneity within the data sets. The ComK-DNA data set was finally classified into 6 3D classes, and the 4 best resolved classes were then subjected to homogenous refinement (final structures in 180 pixel box, 2.18 Å/pixel). The free DNA reconstructions showed limited resolution throughout, therefore we concentrated on classification based on overall curvature. The data set was classified into 5 3D classes, followed by 3D homogenous refinement within classes (120 pixel box size, 3.27 Å/pixel). Two of the classes, which showed the same curvature, were merged during refinement. This resulted in 4 free DNA classes of varying curvatures.
Unfiltered half maps of all classes were subjected to resolution estimate and preprocessing using RELION software[17]. Global resolution was estimated according to the gold-standard FSC=0.143

criteria using the corrected FSC curves (Extended Data Fig. 8-9). Local resolution estimate and filtering was performed using RELION. Attempts to limit the refinements by applying various masks, including focusing on single ComK binding sites (ComK dimers), did not improve the resolution. All ComK-DNA structures featured preferred orientation (Extended Data Fig. 8b), whereas the free DNA structures featured a relatively variable distribution of orientation (Extended Data Fig. 9b). 3D visualization was performed using UCSF Chimera[18] or ChimeraX[19].

We used the same processing procedure to analyze the ComK-*addAB* data set. Following motion correction and CTF estimation, 10,526 micrographs were retained for processing. Automated picking followed by selection of particles by 2D classification yielded 355,612 good particles. However, attempts to perform 3D reconstructions failed because the ComK-*addAB* complex featured severe preferred orientations.

**Classification of ComK-DNA complexes.** To determine the binding stoichiometry of ComK to DNA we slightly modified the classification strategy compared to the one described above, which aimed at achieving well-resolved 3D reconstructions. The initial automatically-picked particle data sets (ComK-*comG* or ComK-*addAB*) were iteratively aligned and classified in 2D, resulting in three clear classes: free DNA, ComK bound to DNA in one box, and ComK bound to DNA in two boxes (Fig. 3b). Exceptionally long DNA class averages appeared in a subset of the class averages, occasionally with three ComK densities. Sub-classification of these elongated DNA classes revealed that they were artifacts due to a mismatch in aligning some of the complexes. Additionally, the current classification included particles, which were discarded before the 3D reconstruction above. This is because the criteria for selecting 2D class averages for 3D reconstruction was stricter with regard to the appearance of fine features in the 2D classes.

**Estimation of the ComK mass in 3D reconstructions.** The volumes associated with ComK densities in the 3D classes were estimated separately for each box, resulting in 8 independent measurements (2 in each of the final 3D classes in Extended Data Fig. 7-8). Volume measurements were done after subtracting the DNA portion from the maps, based on the fitted DNA model, and erasing all density flanking the relevant ComK complex (Fig. 3c). The analysis was performed using UCSF Chimera[18]. Specifically, 20-bp segments, which overlapped with ComK density, were extracted from the DNA model and their fitting within each 3D class was adjusted using rigid-body fitting. The models were converted to density maps using the molmap function in Chimera, applying a resolution of 15 Å, which (for this function) matched the level of details in the 3D classes. The DNA densities were converted to masks by binarizing them at a level that matched the volume of the DNA models (using RELION), and these mask were then used to exclude the DNA density from corresponding ComK density, using the mask function in Chimera. Subsequently, the volume eraser function in Chimera was applied to clear the remaining density flanking ComK densities. The isosurface threshold for calculating the volume of the ComK densities was set based on the diameter of the DNA immediately adjacent to ComK. Volume measurements were converted to mass using a density factor of 1.37 g/cm$^3$ [20].

**Calculation of curvature angles of free and ComK-bound *comG*.** Curvature angles (Fig. 3d,e) were computed from a coarse-grained spline that was determined from the electron density of free DNA and of the ComK-*comG* complex. The electron densities were first binned (4 z-slices per bin for free DNA, 3 z-zlices per bin for ComK-*comG*) and subsequently fit with a Bezier 3D-spline using Mathematica 10.3 (Wolfram). The spline was subsequently divided into segments of $b$ = 0.34 nm length. The resulting coordinates $\mathbf{u}_i$ were used to compute the vector $\mathbf{v}$ of each segment $i$ via $\mathbf{v}_i = \mathbf{u}_{i+1} - \mathbf{u}_i$. The angle between neighboured segments was finally computed according to $\theta_i = \cos^{-1} \mathbf{v}_{i+1}\mathbf{v}_i / b^2$. To calculate the expected bending angle from the decay length of the coupling free energies (Fig. 4c), we used the worm-like chain model that predicts an energetic cost of 1.5 kJ mol$^{-1}$ deg$^{-1}$ for bending a 10 bp DNA duplex[21], which converts to 0.73 pN/deg/bp. With the experimentally obtained value of 0.15 ± 0.07 pN/bp for the 21 bp decay length, we expect a bending angle of (0.15 ± 0.07 )/0.73 = 0.21 ± 0.1 deg.

**DNA model building and refinement.** An idealized 93 bp long B-DNA atomic model was built using Coot[22]. The resulting straight double helix with the *comG* DNA sequence was used as a starting model for cycles of real-space rigid-body refinements and manual molecular fittings to the 3D class 1 (Extended Data Fig. 7) with the whole duplex defined as one group. This results in two idealized models of opposite direction with positional uncertainty along the long axis of the EM map. However, both were compatible with the length and the diameter of the map at an absolute contour level between 0.4 and 0.6. In addition, the undulating shape of the electron density of class 1 corresponds well to the position and spacing of major and minor grooves of the DNA, a feature that was later used to position the double helix in the EM-map. Yet, compared to the straight B-DNA model, the electron density in the ComK-*comG* complex was asymmetrically curved. Subsequent real-space rigid-body refinements using Phenix[23] lead to correlation coefficients (CC) of CC(mask) = 0.30 and CC(box) = 0.45 for the whole model and CC(mask)=0.66 and CC(box)=0.65 for a fragment of 37 bp (from nucleotide 24 to 60). The discrepancy between the correlation coefficients of the whole model and short fragment is due to the intrinsic curvature of the EM map, i.e., the curvature of *comG*. The result indicates that the short fragment was correctly placed. Not surprisingly, identical scores were obtained for the two possible directions of the models related by a 2-fold axis by construction. Starting from the short fragment, we used manual modelling based on expected DNA sequence-related features, in particular for the A-tracts of 4 bp or longer. To this end, we identified crystal structures of duplex-DNA containing sequence motifs that are also present in *comG*. In total, 69 structures with a resolution ≤ 2.5 Å, ligand-free and with standard Watson-Crick base-pairing were checked and 9 of them were selected (Extended Data Table 6).

We started by modifying the initial 37 bp duplex fragment and subsequently extended the building process towards each end in blocks. Each modification of a new block (Extended Data Table 5) was followed by cycles of regularization to ensure connectivity and standard geometry using Coot[22] and real-space refinement with Phenix[23]. The progress was monitored using MolProbity[24]. Importantly, only one of the two opposite directions significantly improved and successfully described the shape of the map, including the shape resulting from narrow minor grooves attributed to the A-tracts. Assuming that two ComKs bind their respective box in a similar manner, we were able to position the double-helix along the long axis of the map with a level of uncertainty of $\pm 1$ base-pairs. The final refinement cycle with Phenix[23] includes restraints on bond lengths and bond angles, on base-pairing H-bonds, planarity, parallelism, and on stacking. Due to the low resolution, the atomic displacement parameters (ADPs or B-factors) and the occupancies were not refined but set to $30.0\text{Å}^2$ and 1.00, respectively. The refined *comG* model includes 170 nucleotides in two chains (85 bp). The refinement statistics are given in the Extended Data Table 7.

**Methods references.**


1 Hamoen, L. W., Van Werkhoven, A. F., Bijlsma, J. J., Dubnau, D. & Venema, G. The competence transcription factor of Bacillus subtilis recognizes short A/T-rich sequences arranged in a unique, flexible pattern along the DNA helix. *Genes & Development* **12**, 1539-1550 (1998).

2 Masoud, R. *et al.* Studying the structural dynamics of bipedal DNA motors with single-molecule fluorescence spectroscopy. *ACS Nano* **6**, 6272-6283 (2012).

3 Dertinger, T. *et al.* Two-focus fluorescence correlation spectroscopy: a new tool for accurate and absolute diffusion measurements. *Chemphyschem* **8**, 433-443, doi:10.1002/cphc.200600638 (2007).

4 Bennett, B. D. *et al.* Absolute metabolite concentrations and implied enzyme active site occupancy in Escherichia coli. *Nat Chem Biol* **5**, 593-599 (2009).

5 Lange, C. & Rudolph, R. Suppression of protein aggregation by L-arginine. *Curr Pharm Biotechnol* **10**, 408-414 (2009).

6 Müller, B. K., Zaychikov, E., Bräuchle, C. & Lamb, D. C. Pulsed interleaved excitation. *Biophys J* **89**, 3508-3522 (2005).

7 Schuler, B., Lipman, E. & Eaton, W. Probing the free-energy surface for protein folding with single-molecule fluorescence spectroscopy. *Nature* **419**, 743-747, doi:10.1038/nature01060 (2002).

8 Hofmann, H. *et al.* Polymer scaling laws of unfolded and intrinsically disordered proteins quantified with single-molecule spectroscopy. *Proc Natl Acad Sci USA* **109**, 16155-16160, doi:10.1073/pnas.1207719109 (2012).

9 Schuler, B., Soranno, A., Hofmann, H. & Nettels, D. Single-Molecule FRET Spectroscopy and the Polymer Physics of Unfolded and Intrinsically Disordered Proteins. *Annu Rev Biophys* **45**, 207–231, doi:10.1146/annurev-biophys-062215-010915 (2016).

10 Schuler, B., Müller-Späth, S., Soranno, A. & Nettels, D. Application of confocal single-molecule FRET to intrinsically disordered proteins. *Methods Mol Biol* **896**, 21-45, doi:10.1007/978-1-4614-3704-8_2 (2012).

11 Benke, S., Nettels, D., Hofmann, H. & Schuler, B. Quantifying kinetics from time series of single-molecule Förster resonance energy transfer efficiency histograms. *Nanotechnology* **28**, 114002, doi:10.1088/1361-6528/aa5abd (2017).

12 Clegg, R. M., Murchie, A. I., Zechel, A. & Lilley, D. M. Observing the helical geometry of double-stranded DNA in solution by fluorescence resonance energy transfer. *Proc Natl Acad Sci U S A* **90**, 2994-2998 (1993).

13 Wozniak, A. K., Schröder, G. F., Grubmüller, H., Seidel, C. A. M. & Oesterhelt, F. Single-molecule FRET measures bends and kinks in DNA. *Proc Natl Acad Sci USA* **105**, 18337-18342 (2008).

14 Mastronarde, D. N. Automated electron microscope tomography using robust prediction of specimen movements. *J Struct Biol* **152**, 36-51 (2005).

15 Punjani, A., Rubinstein, J. L., Fleet, D. J. & Brubaker, M. A. cryoSPARC: algorithms for rapid unsupervised cryo-EM structure determination. *Nat Methods* **14**, 290-296 (2017).

16 Rohou, A. & Grigorieff, N. CTFFIND4: Fast and accurate defocus estimation from electron micrographs. *J Struct Biol* **192**, 216-221 (2015).

17 Zivanov, J. *et al.* New tools for automated high-resolution cryo-EM structure determination in RELION-3. *eLife* **7**, 163 (2018).

18 Pettersen, E. F. *et al.* UCSF Chimera--a visualization system for exploratory research and analysis. *Journal of computational chemistry* **25**, 1605-1612 (2004).

19 Goddard, T. D. *et al.* UCSF ChimeraX: Meeting modern challenges in visualization and analysis. *Protein Sci* **27**, 14-25 (2018).

20 Squire, P. G. & Himmel, M. E. Hydrodynamics and protein hydration. *Arch Biochem Biophys* **196**, 165-177 (1979).

21 Privalov, P. L., Dragan, A. I. & Crane-Robinson, C. The cost of DNA bending. *Trends Biochem Sci* **34**, 464-470 (2009).

22 Emsley, P. & Cowtan, K. Coot: model-building tools for molecular graphics. *Acta Crystallogr D Biol Crystallogr* **60**, 2126-2132 (2004).

23 Liebschner, D. *et al.* Macromolecular structure determination using X-rays, neutrons and electrons: recent developments in Phenix. *Acta Crystallogr D Struct Biol* **75**, 861-877 (2019).

24 Williams, C. J. *et al.* MolProbity: More and better reference data for improved all-atom structure validation. *Protein Sci* **27**, 293-315 (2018).

25 Ruegg, T. L. *et al.* Jungle Express is a versatile repressor system for tight transcriptional control. *Nat Comms* **9**, 3617-3613 (2018).

26 DiGabriele, A. D. & Steitz, T. A. A DNA dodecamer containing an adenine tract crystallizes in a unique lattice and exhibits a new bend. *J Mol Biol* **231**, 1024-1039 (1993).

27 Gewirth, D. T. & Sigler, P. B. The basis for half-site specificity explored through a non-cognate steroid receptor-DNA complex. *Nat Struct Biol* **2**, 386-394 (1995).

28 Bonnet, M. *et al.* The structure of Bradyrhizobium japonicum transcription factor FixK2 unveils sites of DNA binding and oxidation. *J Biol Chem* **288**, 14238-14246 (2013).

29 Rodgers, D. W. & Harrison, S. C. The complex between phage 434 repressor DNA-binding domain and operator site OR3: structural differences between consensus and non-consensus half-sites. *Structure* **1**, 227-240 (1993).

30 Todd, A. K. & Neidle, S. unpublished. (2004).

31 Meijsing, S. H. *et al.* DNA Binding Site Sequence Directs Glucocorticoid Receptor Structure and Activity. *Science* **324**, 407-410 (2009).

32 Rhee, S., Martin, R. G., Rosner, J. L. & Davies, D. R. A novel DNA-binding motif in MarA: the first structure for an AraC family transcriptional activator. *Proc Natl Acad Sci USA* **95**, 10413-10418 (1998).

33 Shevtsov, M. B. *et al.* Structural analysis of DNA binding by C.Csp231I, a member of a novel class of R-M controller proteins regulating gene expression. *Acta Crystallogr D Biol Crystallogr* **71**, 398-407 (2015).

#### Extended Data.

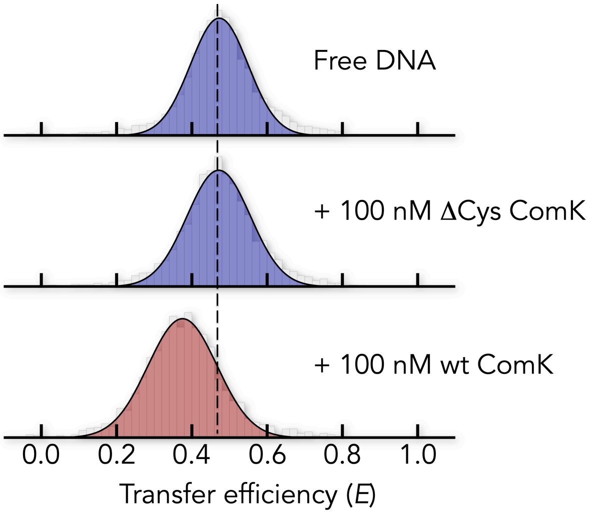


**Extended Data Fig. 1. The cysteine-free variant ΔCys ComK does not bind DNA.** The ComG promoter sequence was labelled at box 2 (Extended Data Table 1). 30 pM DNA (top) was mixed with 100 nM of ΔCys ComK (middle) and wildtype (wt) ComK (bottom). The dashed line is cantered at the FRET histogram of the free DNA.

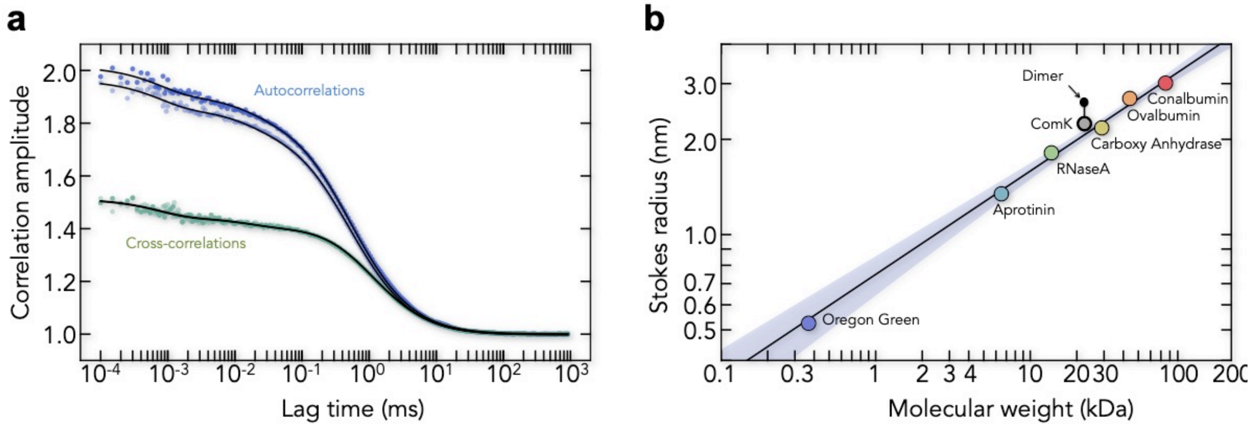


**Extended Data Fig. 2. ComK is a monomer in solution.** (**a**) 2fFCS auto- and cross-correlation functions of singly labelled ComK and fits with a model containing two triplet relaxation processes and one decay for the diffusion of ComK. (**b**) Sokes radius of ComK (grey) in comparison with the Stokes radii of our calibration standards. Based on the measured Stokes radius of ComK, we estimated the radius of a ComK dimer assuming a spherical shape (black dot).

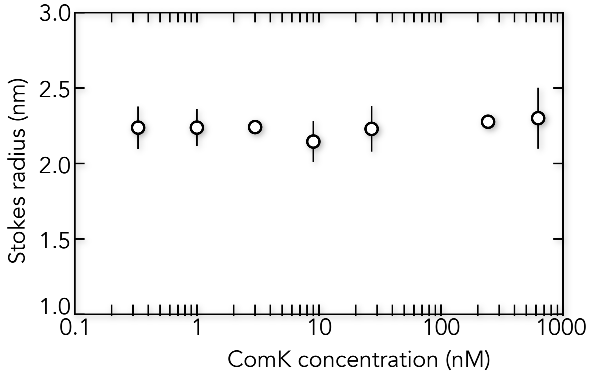


**Extended Data Fig. 3. The dimerization affinity of ComK is low.** The Stokes radius of ComK at increasing concentrations of unlabelled ComK does not increase, indicating that the dissociation constant for homodimerization is > 1 μM.

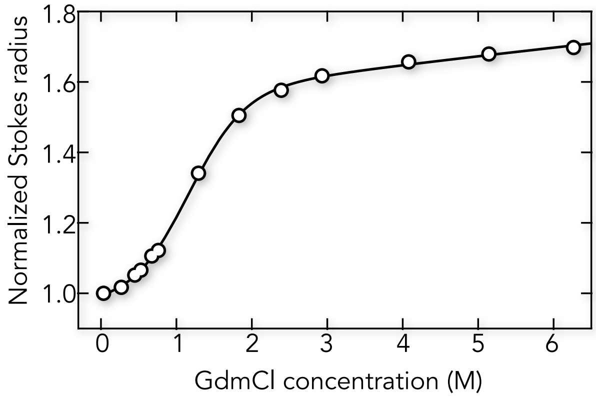


**Extended Data Fig. 4. Thermodynamic stability of ComK.** The Stokes radius relative to the radius in the absence of GdmCl is shown as function of the denaturant (GdmCl) concentration. Solid line is a fit with a two-state folding equilibrium that results in a free energy difference between folded and unfolded ComK of -8.8 ± 0.3 kJ/mol, suggesting that 2.4 ± 0.7 % of the molecules are unfolded under physiological conditions, which explains the strong aggregation tendency of ComK.

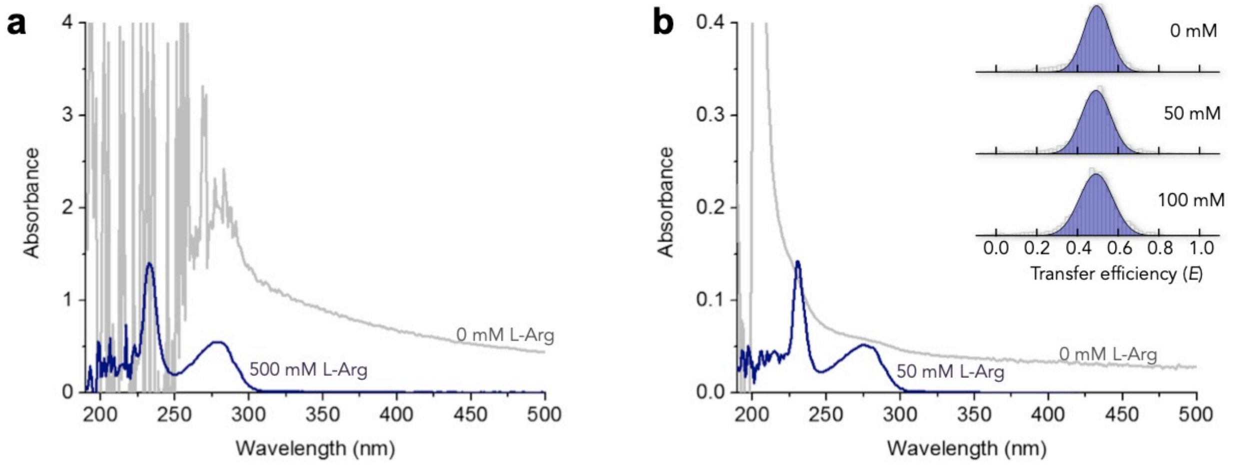


**Extended Data Fig. 5. Impact of L-Arg on ComK and DNA.** Absorbance spectra show that ComK forms aggregates at 22 μM (**a**) and 2 μM (**b**) ComK-concentration (gray lines). However, the wavelength-dependent light-scattering decay is absent in the presence of L-Arg in concentrations of 500 mM (**a**) and 50 mM (**b**). Inset: FRET histograms of labelled *comG* promoter at different L-Arg concentrations (indicated).

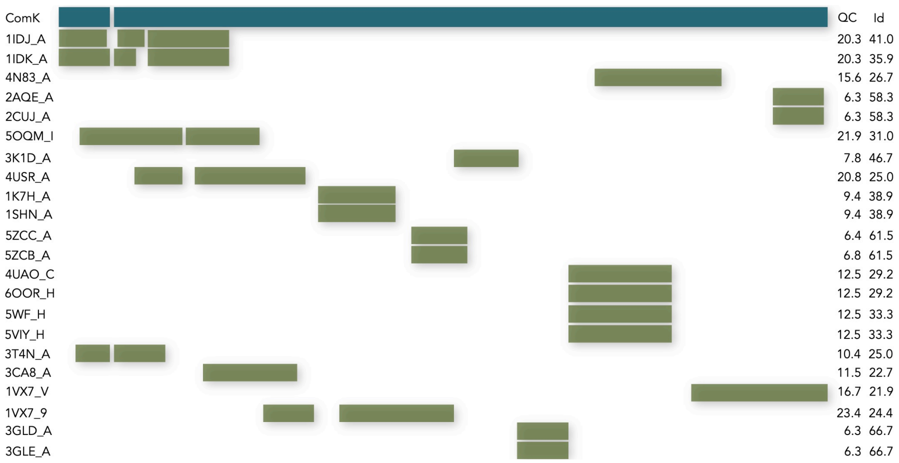


**Extended Data Fig. 6. Multiple sequence alignment of ComK.** The coverage of the ComK sequence (QC in %) and the sequence identities (Id in %) of the covered regions (green bars) are shown in comparison to the ComK sequence (dark cyan).

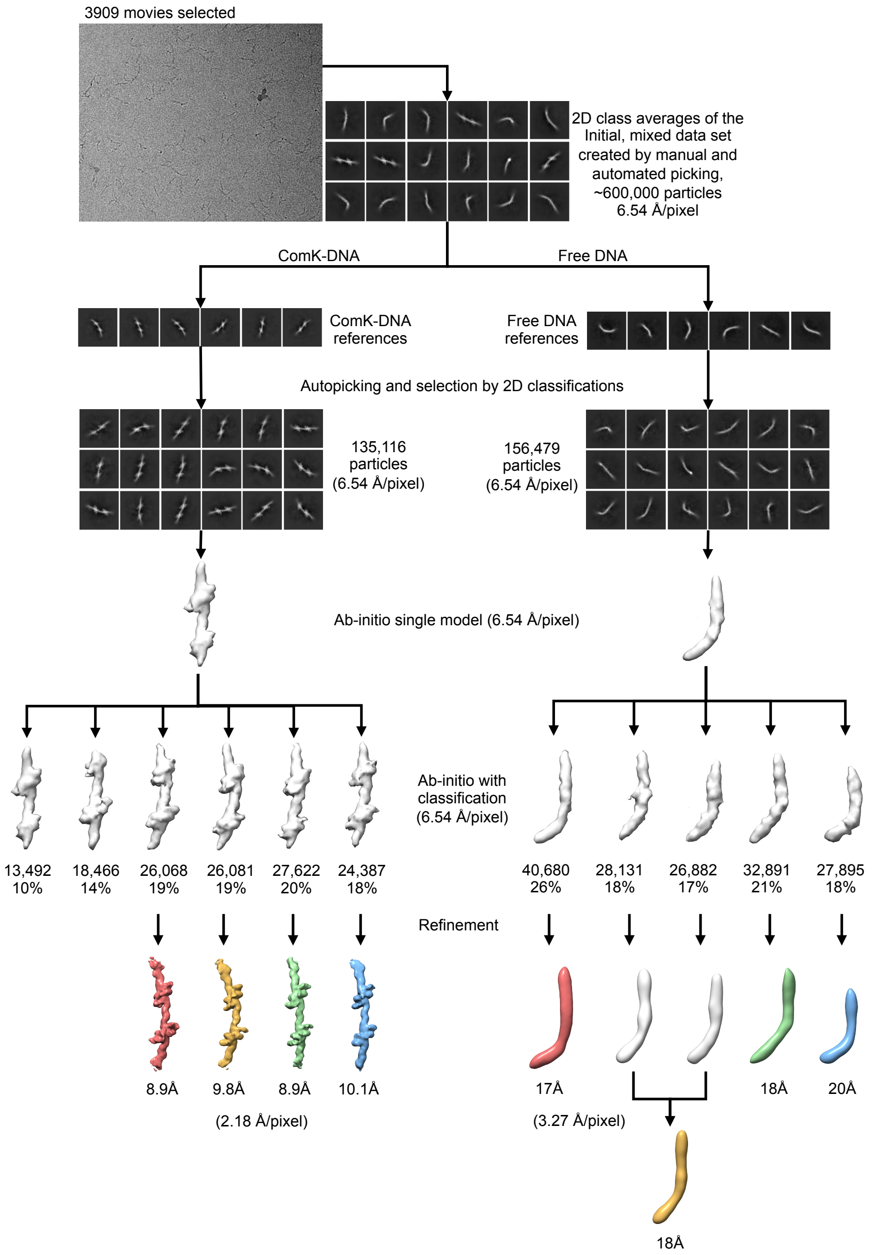


**Extended Data Fig. 7. Scheme for single particle cryo-EM analysis of the ComK-*comG* data set.** The details of the process are described in the supplementary methods section. Briefly, a mixed particle data set was first extracted from the micrographs, containing both free and ComK-bound DNA complexes. 2D class averages of the ComK-DNA or the free DNA were used as references for another round of auto-picking, resulting two separate data sets, ComK-DNA and free DNA. The two data sets were then cleaned by iterative selection of 2D classes, followed by ab-initio 3D reconstruction, 3D classification and refinement of the best 3D classes. Bottom: Classes 1 – 4 (from left to right) for the ComK-DNA complex (left) and free DNA (right).

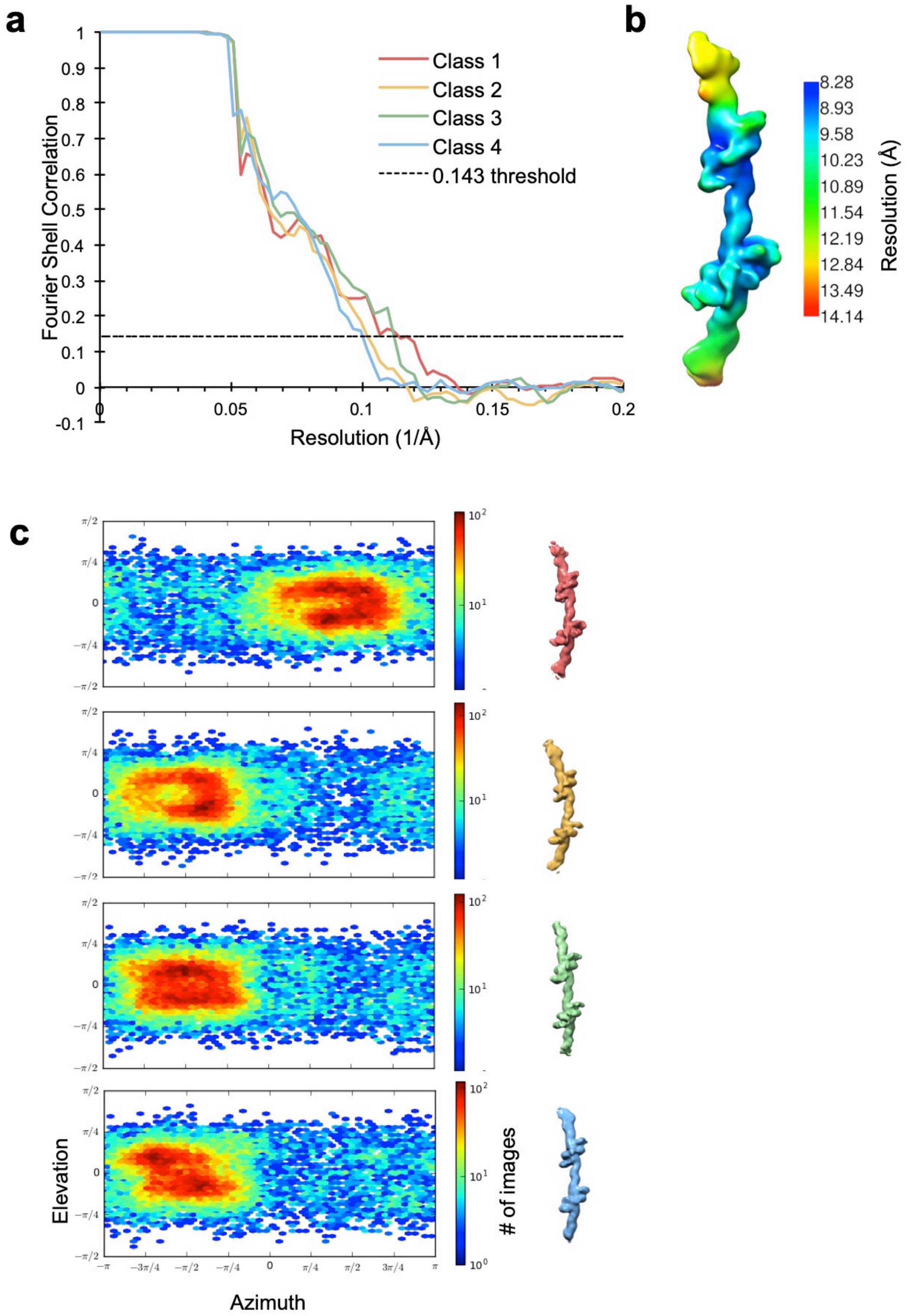


**Extended Data Fig. 8. Analysis of the ComK-*comG* 3D classes.** (**a**) Gold-standard Fourier Shell Correlation (FSC) of the masked, corrected 3D reconstructions (as defined in RELION)[17]. (**b**) Class 1 reconstruction coloured according to local resolution estimate. (**c**) Angular distribution plots for the four 3D classes.

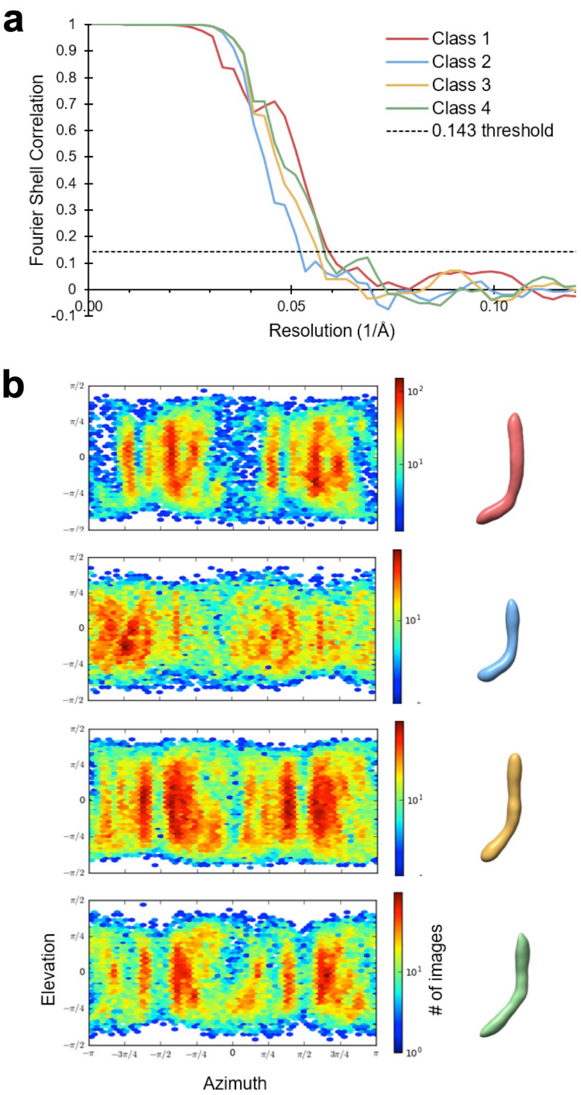


**Extended Data Fig. 9. Analysis of the free *comG* DNA 3D classes.** (**a**) Gold-standard Fourier Shell Correlation (FSC) of the masked, corrected 3D reconstructions (as defined in RELION)[17]. (**b**) Angular distribution plots for the three 3D classes.

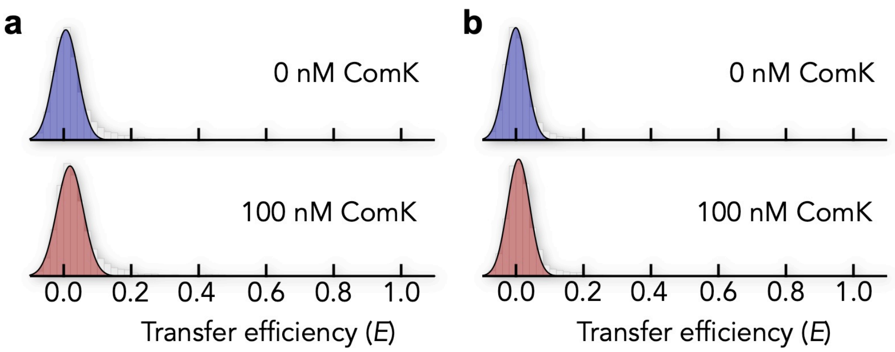


**Extended Data Fig. 10. FRET histograms of the terminally labelled *comG*-promoter.** The acceptor was placed 5' of box 1 whereas the donor was placed 3' of box2. The donor-acceptor sequence separation is 45 bp (a) and 69 bp (b).

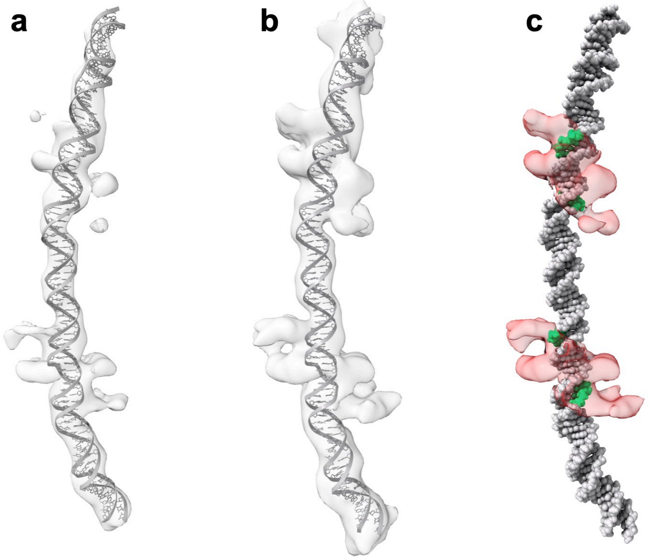


**Extended Data Fig. 11. Flexible fitting of an atomic model of the *comG* promoter DNA into the 3D reconstruction of the ComK-DNA complex.** (**a**, **b**) DNA model and electron density of the complex at two different iso-surface thresholds. At the higher iso-surface threshold (**a**), the ComK density is removed, while the bulk of the DNA density remains. (**c**) Overlay of the atomic model of the *comG* DNA with the electron density of ComK. The DNA density was subtracted for display purposes and the ComK density (red) is shown at an iso-surface threshold to emphasize the DNA binding sites, as in Fig. 3c. The thymine bases at the A-tracts are coloured green.

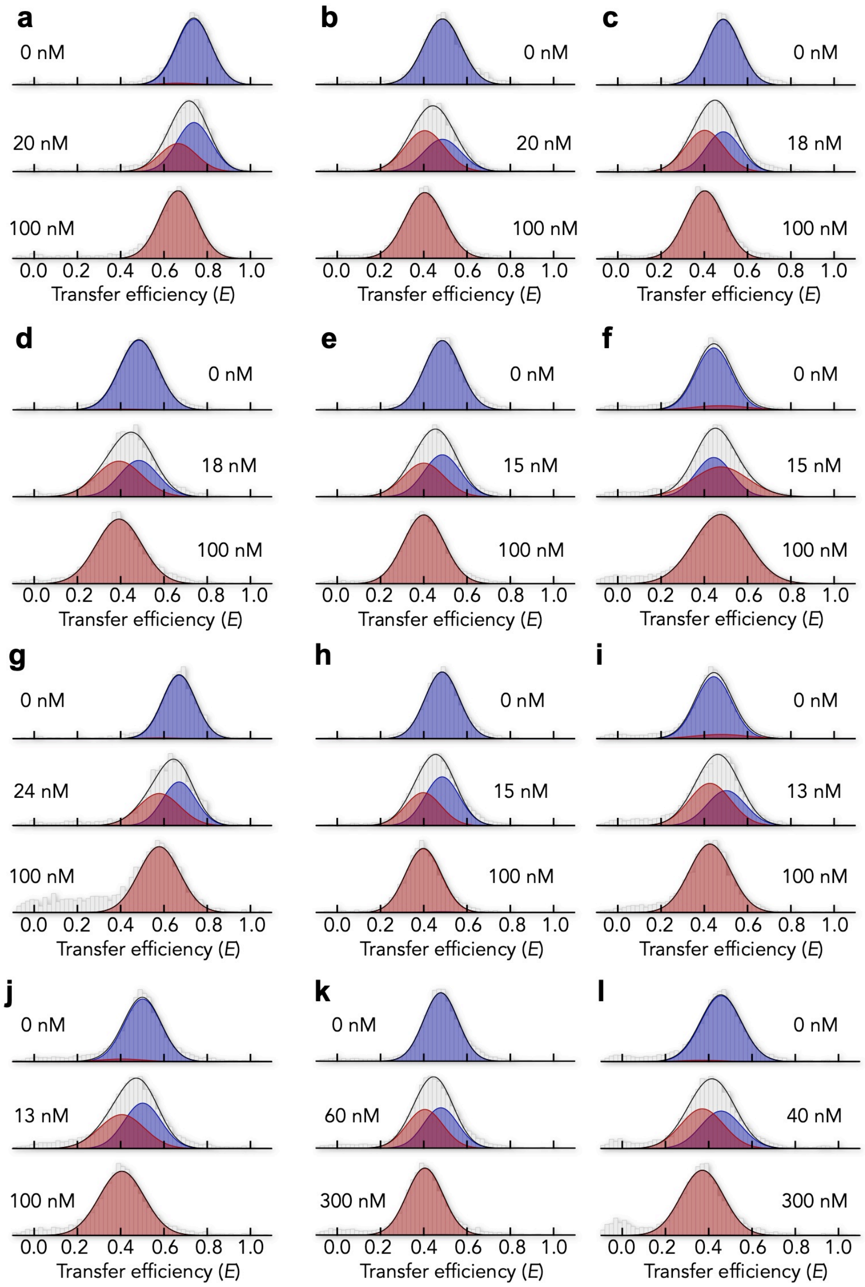


**Extended Data Fig. 12. FRET histograms of all constructs.** Histograms are shown for the promoters *addAB* with labelled box 1 (**a**), *addAB* with labelled box 2 (**b**), *comG* with 55% GC content in the spacer (**c**), *comG* with 72% GC content in the spacer (**d**), *comG* with mismatch in the spacer (**e**), *comG* with a nick in the spacer (**f**), *comK* promoter (**g**), artificial promoter with 8 bp spacers labelled in box 2 and containing 2 boxes (**h**), 3 boxes (**i**), and 4 boxes (**j**), the isolated box 1 of *comG* (**k**), and the isolated box 2 of *comG* (**l**). Unless otherwise state, box 1 is labelled. The concentration of ComK is indicated.

**Extended Data Table 1. List of labelled DNA for smFRET experiments that monitor the curvature in the *comG* promoter.** The two AT boxes are shaded in cyan. Donor and acceptor attachment sites are shown in green and red, respectively. Duplexes were obtained from combinations of the three Watson strands (W) with the four Crick strands (C), thus resulting in the 11 constructs shown in Fig. 2b.

| *comG* | Length | Sequence |
|---|---|---|
| **W71** | 84 | CAGTTGAAAGTCTTTTTTCTTGCCAGAAAGAATTGGTTTTTCAGCATATAACATCTCACAAAATCACGTTTTCCCTGTTTGATT |
| ***W64*** | 84 | CAGTTGAAAGTCTTTTTTCTTGCCAGAAAGAATTGGTTTTTCAGCATATAACATCTCACAAAATCACGTTTTCCCTGTTTGATT |
| ***W41*** | 84 | CAGTTGAAAGTCTTTTTTCTTGCCAGAAAGAATTGGTTTTTCAGCATATAACATCTCACAAAATCACGTTTTCCCTGTTTGATT |
| ***C66*** | 84 | AATCAAACAGGGAAAACGTGATTTTGTGAGATGTTATATGCTGAAAAACCAATTCTTTCTGGCAAGAAAAAAGACTTTCAACTG |
| ***C62*** | 84 | AATCAAACAGGGAAAACGTGATTTTGTGAGATGTTATATGCTGAAAAACCAATTCTTTCTGGCAAGAAAAAAGACTTTCAACTG |
| ***C58*** | 84 | AATCAAACAGGGAAAACGTGATTTTGTGAGATGTTATATGCTGAAAAACCAATTCTTTCTGGCAAGAAAAAAGACTTTCAACTG |
| ***C43*** | 84 | AATCAAACAGGGAAAACGTGATTTTGTGAGATGTTATATGCTGAAAAACCAATTCTTTCTGGCAAGAAAAAAGACTTTCAACTG |
| ***C32*** | 84 | AATCAAACAGGGAAAACGTGATTTTGTGAGATGTTATATGCTGAAAAACCAATTCTTTCTGGCAAGAAAAAAGACTTTCAACTG |

**Extended Data Table 2. List of labelled DNA for smFRET ComK-binding experiments.** The two AT boxes are highlighted in cyan. Donor and acceptor attachment sites are shown in green and red, respectively. For the constructs of the isolated boxes, we added a flanking region (15-17 bp) before and after each box with GC pairs at the termini to prevent helix fraying.

| Promoter | Length | Sequence |
|---|---|---|
| ***comG*: box 1 labelled** | 93 | CAGTTGAAAGTCTTTTTTCTTGCCAGAAAGAATTGGTTTTTCAGCATATAACATCTCACAAAATCACGTTTTCCCTGTTTGATTACCTTTTCT<br>GTCAACTTTCAGAAAAAAGAACGGTCTTTCTTAACCAAAAAGTCGTATATTGTAGAGTGTTTTAGTGCAAAAGGGACAAACTAATGGAAAAGA |
| ***comG*: spacer labelled** | | CAGTTGAAAGTCTTTTTTCTTGCCAGAAAGAATTGGTTTTTCAGCATATAACATCTCACAAAATCACGTTTTCCCTGTTTGATTACCTTTTCT<br>GTCAACTTTCAGAAAAAAGAACGGTCTTTCTTAACCAAAAAGTCGTATATTGTAGAGTGTTTTAGTGCAAAAGGGACAAACTAATGGAAAAGA |
| ***comG*: box 2 labelled** | 93 | CAGTTGAAAGTCTTTTTTCTTGCCAGAAAGAATTGGTTTTTCAGCATATAACATCTCACAAAATCACGTTTTCCCTGTTTGATTACCTTTTCT<br>GTCAACTTTCAGAAAAAAGAACGGTCTTTCTTAACCAAAAAGTCGTATATTGTAGAGTGTTTTAGTGCAAAAGGGACAAACTAATGGAAAAGA |
| ***comG*: isolated box 1 labelled** | 46 | CTTTTTTCTTGCCAGAAAGAATTGGTTTTTCAGCATATAACATCTC<br>GAAAAAAGAACGGTCTTTCTTAACCAAAAAGTCGTATATTGTAGAG |
| ***comG*: isolated box 2 labelled** | 43 | CATATAACATCTCACAAAATCACGTTTTCCCTGTTTGATTACC<br>GTATATTGTAGAGTGTTTTAGTGCAAAAGGGACAAACTAATGG |
| ***comG* GC-55%: box 1 labelled** | 93 | CAGTTGAAAGTCTTTTTTCTTGCCAGAAAGAATTGGTTTTTCAGCAGCTAACAGCTCACAAAATCACGTTTTCCCTGTTTGATTACCTTTTCT<br>GTCAACTTTCAGAAAAAAGAACGGTCTTTCTTAACCAAAAAGTCGTCGATTGTCGAGTGTTTTAGTGCAAAAGGGACAAACTAATGGAAAAGA |
| ***comG* GC-72%: box 1 labelled** | 93 | CAGTTGAAAGTCTTTTTTCTTGCCAGAAAGAATTGGTTTTTCGGCCGCTAACGGCTCACAAAATCACGTTTTCCCTGTTTGATTACCTTTTCT<br>GTCAACTTTCAGAAAAAAGAACGGTCTTTCTTAACCAAAAAGCCGGCGATTGCCGAGTGTTTTAGTGCAAAAGGGACAAACTAATGGAAAAGA |
| ***comG* Mismatch: box 1 labelled** | 93 | CAGTTGAAAGTCTTTTTTCTTGCCAGAAAGAATTGGTTTTTCAGCAATATACATCTCACAAAATCACGTTTTCCCTGTTTGATTACCTTTTCT<br>GTCAACTTTCAGAAAAAAGAACGGTCTTTCTTAACCAAAAAGTCGTATATTGTAGAGTGTTTTAGTGCAAAAGGGACAAACTAATGGAAAAGA |
| ***comG* Nicked: box 1 labelled** | 50<br>43<br>93 | CAGTTGAAAGTCTTTTTTCTTGCCAGAAAGAATTGGTTTTTCAGCATATA<br>ACATCTCACAAAATCACGTTTTCCCTGTTTGATTACCTTTTCT<br>GTCAACTTTCAGAAAAAAGAACGGTCTTTCTTAACCAACAAGTCGTATATTGTAGAGTGTTTTAGTGCACAAGGGACAAACTAATGGAAAAGA |
| ***comK*: box 1 labelled** | 78 | CTTATGAAAGTAAAATCGGTTTATTACTAGTCATTTAGTACCATTAAATATCATTAAAAGATGATTTTATCTTAAATG<br>GAATACTTTCATTTTAGCCAAATAATGATCAGTAAATCATGGTAATTTATAGTAATTTTCTACTAAAATAGAATTTAC |
| ***addAB1*: box 1 labelled** | 57 | GATGTGCGGAGATAATCAGCTTTTTATATGTGAAAAGGCCGTTTTTACCAATAGATC<br>CTACACGCCTCTATTAGTCGAAAAATATACACTTTTCCGGCAAAAATGGTTATCTAG |
| ***addAB2*: box 2 labelled** | 57 | GATGTGCGGAGATAATCAGCTTTTTATATGTGAAAAGGCCGTTTTTACCAATAGATC<br>CTACACGCCTCTATTAGTCGAAAAATATACACTTTTCCGGCAAAAATGGTTATCTAG |
| ***addAB 2 boxes*** | 57 | GATGTGCGGAGAAAAGGCCGTTTTTATATGTGAAAAGGCCGTTTTTACCAATAGATC<br>CTACACGCCTCTTTTCCGGCAAAAATATACACTTTTCCGGCAAAAATGGTTATCTAG |
| ***addAB 3 boxes*** | 78 | GATGTGCGGAGAAAAGGCCGTTTTTATATGTGAAAAGGCCGTTTTTATATGTGAAAAGGCCGTTTTTACCAATAGATC<br>CTACACGCCTCTTTTCCGGCAAAAATATACACTTTTCCGGCAAAAATATACACTTTTCCGGCAAAAATGGTTATCTAG |
| ***addAB 4 boxes*** | 99 | GATGTGCGGAGAAAAGGCCGTTTTTATATGTGAAAAGGCCGTTTTTATATGTGAAAAGGCCGTTTTTATATGTGAAAAGGCCGTTTTTACCAA<br>TAGATC<br>CTACACGCCTCTTTTCCGGCAAAAATATACACTTTTCCGGCAAAAATATACACTTTTCCGGCAAAAATATACACTTTTCCGGCAAAAATGGTT<br>ATCTAG |

**Extended Data Table 3. Amino acid sequences of the ComK constructs used in this study.** The codon optimized construct contained a $His_6$-tag (red) and an HRV3C cleavage site (cyan) used for all smFRET and cryo-EM experiments (top). For determination of the oligomerization state using 2fFCS, we also created a construct with an additional cysteine at the N-terminus (bottom). Vertical line indicates the HRV3C cleavage position.

**ComK amino acid sequence**

MGSSHHHHHHSGSGSAGLEVLFQ|GPGMSQKTDAPLESYEVNGATIAVLPEEIDGKICSKIIEKDCVFYVNMKPLQIVDRSCRFFG
SSYAGRKAGTYEVTKISHKPPIMVDPSNQIFLFPTLSSTRPQCGWISHVHVKEFKATEFDDTEVTFSNGKTMELPISYNSFENQVY
RTAWLRTKFQDRIDHRVPKRQEFMLYPKEERTKMIYDFILRELGERY

MGSSHHHHHHSGSGSAGLEVLFQ|GPCMSQKTDAPLESYEVNGATIAVLPEEIDGKICSKIIEKDCVFYVNMKPLQIVDRSCRFFG
SSYAGRKAGTYEVTKISHKPPIMVDPSNQIFLFPTLSSTRPQCGWISHVHVKEFKATEFDDTEVTFSNGKTMELPISYNSFENQVY
RTAWLRTKFQDRIDHRVPKRQEFMLYPKEERTKMIYDFILRELGERY

**Extended Data Table 4. Fitting parameters obtained with the mechanistic binding model and with the Hill equation** Errors result from at least two independent experiments. To increase the robustness of the fit parameters in our mechanistic binding model, we fit the constructs in which we probed both boxes (*addAB* and *comG*) in global manner with the same parameter for both constructs. Fixed parameters are indicated by (*) and global fit parameters are indicated by (#).

| Promoter | $\Delta g_K$ ($k_BT$) | $-\Delta g_\sigma$ ($k_BT$) | $-\Delta g_J$ ($k_BT$) | $K_{Hill}$ (nM) | $n$ |
|---|---|---|---|---|---|
| *comG*: box 1 labelled | 7.0 ± 0.6# | 4.0* | 4.5 ± 1.2# | 15 ± 1 | 3.55 ± 0.12 |
| *comG*: box 2 labelled | 7.0 ± 0.6# | 4.0* | 4.5 ± 1.2# | 16 ± 1 | 3.44 ± 0.27 |
| *comG*: isolated box 1 labelled | 6.1 ± 0.7 | 4.0 ± 0.7# | n.d. | 73 ± 21 | 1.76 ± 0.02 |
| *comG*: isolated box 2 labelled | 5.7 ± 0.7 | 4.0 ± 0.7# | n.d. | 48 ± 14 | 1.59 ± 0.09 |
| *comG* GC-55%: box 1 labelled | 6.6 ± 0.7 | 4.0* | 3.7 ± 1.2 | 16 ± 1 | 2.97 ± 0.28 |
| *comG* GC-72%: box 1 labelled | 6.5 ± 0.5 | 4.0* | 3.5 ± 0.9 | 17 ± 1 | 3.05 ± 0.08 |
| *comG* Mismatch: box 1 labelled | 6.1 ± 0.1 | 4.0* | 3.1 ± 0.1 | 14 ± 1 | 2.93 ± 0.28 |
| *comG* Nicked: box 1 labelled | 4.2 ± 0.2 | 4.0* | 0.2 ± 0.1 | 19 ± 8 | 1.82 ± 0.12 |
| *comK*: box 1 labelled | 5.8 ± 0.2 | 4.0* | 1.9 ± 0.1 | 21 ± 2 | 2.28 ± 0.16 |
| *addAB1*: box 1 labelled | 8.0 ± 0.7# | 4.0* | 5.8 ± 0.9# | 25 ± 6 | 4.34 ± 0.32 |
| *addAB2*: box 2 labelled | 8.0 ± 0.7# | 4.0* | 5.8 ± 0.9# | 24 ± 7 | 3.77 ± 0.45 |

**Extended Data Table 5. DNA sequences used for gel retardation assay.** The two AT boxes are highlighted in cyan. Only the forward strand is shown. The 18 bp spacer of the ComG promoter, was modified such that the spacer length was truncated or extended. To keep a uniform length for all dsDNA, the sequence length spanning the two AT boxes was altered, based on the genomic sequence, keeping the termini with a GC pair.

| Promoter | Length | Sequence |
|---|---|---|
| **Spacer: 4 bp** | 89 | GCAGTTGAAAGTCTTTTTTCTTGCCAGAAAGAATTGGTTTTTCAGCAAAATCACGTTTTCCCTGTTTGATTACCTTTTCTTCTTTTTCG |
| **Spacer: 6 bp** | 89 | CAGTTGAAAGTCTTTTTTCTTGCCAGAAAGAATTGGTTTTTCAGCATAAAATCACGTTTTCCCTGTTTGATTACCTTTTCTTCTTTTTC |
| **Spacer: 8 bp** | 89 | CGTTGAAAGTCTTTTTTCTTGCCAGAAAGAATTGGTTTTTCAGCATATAAAATCACGTTTTCCCTGTTTGATTACCTTTTCTTCTTTTG |
| **Spacer: 10 bp** | 89 | GTTGAAAGTCTTTTTTCTTGCCAGAAAGAATTGGTTTTTCAGCATATAAAAAATCACGTTTTCCCTGTTTGATTACCTTTTCTTCTTTG |
| **Spacer: 12 bp** | 89 | CTGAAAGTCTTTTTTCTTGCCAGAAAGAATTGGTTTTTCAGCATATAACAAAAATCACGTTTTCCCTGTTTGATTACCTTTTCTTCTTG |
| **Spacer: 14 bp** | 89 | CGAAAGTCTTTTTTCTTGCCAGAAAGAATTGGTTTTTCAGCATATAACATCAAAATCACGTTTTCCCTGTTTGATTACCTTTTCTTCTG |
| **Spacer: 16 bp** | 89 | GAAAGTCTTTTTTCTTGCCAGAAAGAATTGGTTTTTCAGCATATAACATCTCAAAATCACGTTTTCCCTGTTTGATTACCTTTTCTTCG |
| **Spacer: 18 bp** | 89 | CAAGTCTTTTTTCTTGCCAGAAAGAATTGGTTTTTCAGCATATAACATCTCACAAAATCACGTTTTCCCTGTTTGATTACCTTTTCTTC |
| **Spacer: 20 bp** | 89 | CAGTCTTTTTTCTTGCCAGAAAGAATTGGTTTTTCAGCATATAACATCTCACCAAAAATCACGTTTTCCCTGTTTGATTACCTTTTCTG |
| **Spacer: 22 bp** | 89 | CGTCTTTTTTCTTGCCAGAAAGAATTGGTTTTTCAGCATATAACATCTCACCAGCAAAATCACGTTTTCCCTGTTTGATTACCTTTTCG |
| **Spacer: 22 bp** | 89 | GTCTTTTTTCTTGCCAGAAAGAATTGGTTTTTCAGCATATAACATCTCACCAGCATAAAATCACGTTTTCCCTGTTTGATTACCTTTTC |

**Extended Data Table 6. PDB files used to model specific sequence motifs (white letters).**

a b c d e f g h i

**5′-CAGTTGAAAGTCTTTTTTCTTGCCAGAAAGAATTGGTTTTTCAGCATATAACATCTCACAAAATCACGTTTTCCCTGTTTGATTACCTTTTCT -3′**

| | PDB ID | Resolution (Å) | Sequence[2] | Reference |
|---|---|---|---|---|
| **a**[1] | 5VL9 | 2.2 | GAAAGTt- | 25 |
| **b** | 1D89 | 2.3 | -gTTTTTTCg- | 26 |
| **c** | 1LAT | 1.9 | -tCCAGAAc- | 27 |
| **d** | 4I2O | 1.8 | -gGAATTGa- | 28 |
| **e** | 1PER | 2.5 | -aGTTTTTCt- | 29 |
| **f** | 1LP7 | 2.4 | -cATATAa- | 30 |
| **g** | 3G9M | 1.6 | -aCAAAATg- | 31 |
| **h** | 1BL0 | 2.3 | -cGTTTTg- | 32 |
| **i** | 4JCX | 2.3 | -gTTTTCc- | 33 |

[1]Position of the segment in the 93bp *comG* sequence.
[2]Lowercases indicate flanking bases not added to the model.

**Extended Data Table 7. Refinement statistics.**

| Pseudo space group | *P*1 |
|---|---|
| Box | |
| Lengths (Å) | 63.2, 50.1, 289. 9 |
| Angles (°) | 90.0, 90.0, 90.0 |
| CC (mask) | 0.61 |
| CC(box) | 0.69 |
| CC(peaks) | 0.37 |
| CC(volume) | 0.54 |
| Number of chains | 2 |
| Number of residues | 170 |
| Number of atoms | 3479 |
| Bonds (RMSD) | |
| Bond (Å) (> 4$\sigma$) | 0.009 (0) |
| Angle (°) (> 4$\sigma$) | 1.214 (74) |
| Clashes | 11 |
